# IGF-1 IMPACTS NEOCORTICAL INTERNEURON CONNECTIVITY IN EPILEPTIC SPASM GENERATION AND RESOLUTION

**DOI:** 10.1101/2024.10.13.618035

**Authors:** Carlos J. Ballester-Rosado, John T. Le, Trang T. Lam, Anne E. Anderson, James D. Frost, John W. Swann

**Affiliations:** The Cain Foundation Laboratories, the Jan and Dan Duncan Neurological Research Institute, Texas Children’s Hospital, Houston, Texas, USA; Department of Pediatrics, Baylor College of Medicine, Houston, Texas, USA; Department of Neuroscience, Baylor College of Medicine, Houston, Texas, USA; Department of Neurology, Baylor College of Medicine, Houston, Texas, USA

**Keywords:** Infantile Spasms, IESS, dysmaturation, autocrine signaling, interneurons

## Abstract

Little is known about the mechanisms that generate epileptic spasms following perinatal brain injury. Recent studies have implicated reduced levels of Insulin-like Growth Factor 1 (IGF-1) in these patients’ brains. Other studies have reported low levels of the inhibitory neurotransmitter, GABA. In the TTX brain injury model of epileptic spasms, we undertook experiments to evaluate the impact of IGF-1 deficiencies on neocortical interneurons and their role in spasms. Quantitative immunohistochemical analyses revealed that neocortical interneurons that express glutamic acid decarboxylase, parvalbumin, or synaptotagmin 2 co-express IGF-1. In epileptic rats, expression of these three interneuron markers were reduced in the neocortex. IGF-1 expression was also reduced, but surprisingly this loss was confined to interneurons. Interneuron connectivity was reduced in tandem with IGF-1 deficiencies. Similar changes were observed in surgically resected neocortex from infantile epileptic spasms syndrome (IESS) patients. To evaluate the impact of IGF-1 deficiencies on interneuron development, IGF-1R levels were reduced in the neocortex of neonatal conditional IGF-1R knock out mice by viral injections. Four weeks later, this experimental maneuver resulted in similar reductions in interneuron connectivity. Treatment with the IGF-1 derived tripeptide, (1–3)IGF-1, abolished epileptic spasms in most animals, rescued interneuron connectivity, and restored neocortical levels of IGF-1. Our results implicate interneuron IGF-1 deficiencies, possibly impaired autocrine IGF-1 signaling and a resultant interneuron dysmaturation in epileptic spasm generation. By restoring IGF-1 levels, (1–3)IGF-1 likely suppresses spasms by rescuing interneuron connectivity. Results point to (1–3)IGF-1 and its analogues as potential novel disease-modifying therapies for this neurodevelopmental disorder.

---

Epileptic spasms is a severe type of epilepsy occurring almost exclusively during infancy and early childhood (1, 2). In infants, it is now called infantile epileptic spasms syndrome (IESS). The seizures are brief flexion and/or extension contractions of the trunk or limbs. With each spasm, a concurrent ictal event is seen in EEG recordings (3) and between spasms a chaotic pattern called hypsarrhythmia is commonly recorded (4). Adrenocorticotropic hormone (ACTH), corticosteroids and vigabatrin can eliminate spasms in 50%, 40% and 30% of infants, respectively (5). However, even after successful treatment, long-term outcomes are poor (6) - especially if treatment is delayed and there is an underlying brain abnormality (7). Thus, current therapies are not considered disease-modifying.

The mechanisms underlying epileptic spasms, hypsarrhythmia, and their neurobehavioral comorbidities are not fully understood (8–10). Over 200 clinically disparate conditions are associated with IESS (6). There is also a heterogeneous list of single gene mutations that are thought to be causative (11). Among them is *Aristaless*-related homeobox gene (*ARX*). Missense mutations of *ARX* or expansion of a polyalanine tract commonly leads to X-linked infantile spasms syndrome (12, 13) and an impaired migration of interneurons into the neocortex (14). Such observations led to a proposed interneuronopathy hypothesis for IESS (15). More recently, several animal models of IESS have reported deficits in inhibitory interneuron populations in the neocortex and altered inhibitory synaptic transmission (16–20).

Beyond genetic etiologies, three studies have reported low levels of GABA in the CSF of IESS patients (21–23). In one study, decreased GABA levels were only observed in patients with a known etiology (“symptomatic” cases) (23). Similarly, Riikonen and colleagues have reported decreased levels of the Insulin-like Growth Factor 1 (IGF-1) in the CSF of children with symptomatic spasms (24). A loss of IGF-1 in the developing brain could have significant consequences, since numerous studies in transgenic mouse models have shown that the elimination of IGF1 leads to a decrease in brain size and overexpression has the opposite effect (25). For instance, in nestin-driven IGF-1 overexpression mice, neocortical weight is increased 29% by P12 (26). On the other hand, nestin-driven elimination of the receptor for IGF-1 (IGF-1R) results in death soon after birth but in heterozygotes a 40% reduction in brain weight is observed (27). Interestingly, IGF-1 overexpression has also been shown to increase synapse number that could not be explained by a concomitant increase in neuron number - highlighting its role in synaptogenesis (28). However, the role of IGF-1 in GABAergic neocortical interneuron development and circuit formation has not been evaluated.

In experiments reported here, we explored abnormalities in neocortical interneurons and the influence IGF-1 may have on them in the TTX animal model of IESS. In these animals, epileptic spasms are induced in rats by infusion of TTX into the cortex beginning in infancy (29). Within one week, animals begin to display epileptic spasms that have neurophysiological and behavioral features and pharmacological sensitivities equivalent to those of humans (29–31). Previously, we reported a widespread loss of IGF-1 in the neocortex of animals with spasms (32). Similar alterations in IGF-1 expression were observed in surgically resected neocortices from symptomatic IESS patients. Treatment with (1–3)IGF-1, which was found to act through the receptor for IGF-1, abolished spasms in the majority of animals. Here we extend that work and report that the loss of IGF-1 in cortex of epileptic animals is largely limited to interneurons and is associated with a reduction in neocortical interneuron connectivity. In animals in which (1–3)IGF-1 abolished spasms, GABAergic nerve terminal networks were rescued and IGF-1 levels restored.

## Methods

### Animals

Wistar rats were obtained from Envigo. IGF1R^lox^ mice were obtained from the Jackson Laboratory (B6;129-IGF1R^tm2Arge^/J) and C57Bl/6J mice from the Baylor College of Medicine CCM Production Colony. All animals were used in a 1:1 male to female ratio. For the IGFR^lox^ mice, exon 3 of the *IGF-1R* gene is flanked by *loxP* sites (33). All procedures were approved by the Baylor College of Medicine IACUC and in keeping with NIH guidelines.

### Neocortical TTX infusions

TTX (Alomone) was chronically infused into the cortex as previously described (29). An osmotic minipump (Alzet Model 2004) containing 12 µM TTX in sterile saline was implanted under the skin of 12-day-old rats. Saline-infused rats and uninfused littermates served as controls. Tubing connected pumps to cannulas implanted into the right motor/sensory cortices.

### Long-term video/EEG recordings

EEGs with video were recorded continuously, 24 hours a day, using methods previously described (34). On P23, electrodes constructed from Teflon-coated stainless-steel wire were implanted 1.0 mm below the cortical surface at six sites in the neocortex. Reference electrodes and isolated system grounds were placed in the cerebellum. Recordings were made on the Nicolet/Viasys instrument. One of us (JDF) read and interpreted recordings and was blinded to experimental treatments.

### Drug Treatments

(1–3)IGF-1 (Bachem) was given daily for three weeks. (1–3)IGF-1 has good oral bioavailability (∼50% compared to i.p.), so we treated rats with 40 mg/kg/day by gavage. Recombinant human IGF-1 (rhIGF-1) was obtained from PeproTech.

### AAV injections

We eliminated IGF-1R from neurons in the neocortex as previously described (32). Under hypothermia-induced anesthesia, we unilaterally injected AAV8-EF1α-mCherry-IRES-Cre (Addgene titer 1.0×10^13^ GC/mL) into the sensory cortex of one-to-three-day-old mice born from IGF1R ^lox^ homozygous crossings. Littermates injected with AAV8-EF1α-EGFP (Addgene titer 1.0×10^13^ GC/mL) served as controls. Two hundred nL of the AAV solution were injected at a rate of 20nL/sec and a depth of 400 μm below the surface of the scalp.

### Immunohistochemistry and Image Processing

Rats and mice were perfused with 4% paraformaldehyde. Brains were vibratome sectioned (100 μm). Sections were permeabilized with 0.3% Triton X-100, blocked with normal goat serum, and incubated with primary antibodies diluted in PBST-1% normal goat serum at 4°C overnight. The following day, sections were incubated with secondary antibodies conjugated with Alexa Fluor 488, 594, or 647 (1:500; Life Technologies). Confocal images were acquired with a Zeiss 880/Airyscan system. Z stacks consisted of 14 confocal slices collected at 0.75µm intervals. All immunohistochemistry processing of tissue sections and confocal imaging were batch processed to ensure that valid comparisons could be made between animals and across experimental groups.

The extent of immunoreactivity was quantified by the widely used Imaris surface rendering algorithm which produces three-dimensional surface reconstruction of immunofluorescent cells (35–39). Reconstructed surfaces were used to quantify the volume of tissue occupied by immunolabeling. The threshold for detecting immunoreactivity was set just above background fluorescence and the same threshold was used for all samples analyzed in each experiment. In some instances, the algorithm did not detect very small weakly stained structures that were close to the background. However, since the same settings were used across sections from all experimental and control animals, any underestimation of immunoreactivity would be the same across experimental groups and would not be expected to impact experimental outcomes. We carefully examined each dataset to confirm the reconstructed images faithfully represented the original immunofluorescence.

To quantify protein colocalization, the widely employed “Coloc” algorithm in Imaris was used (39–44). Specifically, glutamic acid decarboxylase (GAD) or parvalbumin (PV)-positive neurons were reconstructed and the resulting image filtered using a masking tool to only display GAD- or PV-positive cell bodies. The Imaris colocalization algorithm was then used to quantify the percentage of GAD- or PV-positive voxels that co-registered with IGF-1 immunopositivity. For comparison, we processed separate histological sections from the same animals but without the IGF-1 primary antibody. To quantify the colocalization of synaptotagmin 2 with PV and IGF-1 in nerve terminals, the same methods were used but the cell bodies were masked to isolate networks of presynaptic terminals. To compute the density of interneurons, confocal stacks of fluorescent images were imported into Neurolucida (MicroBrightField). Individual interneuron cell bodies were marked, the area analyzed outlined on the computer monitor and interneuron density computed. The investigator carrying out these analyses was blinded to experimental conditions.

Primary antibodies used: rabbit anti-IGF1 (1:400; Abcam AB40657); rabbit anti-IGF1 (1:200; Abcam AB9572); mouse anti-GAD67 (1:500; EMD Millipore MAB5406); rabbit anti-GAD65/67 (1:2000; Abcam AB183999); guinea pig anti-parvalbumin (1:4000; Synaptic Systems 195004); rabbit anti-parvalbumin (1:2000; Abcam AB11427); mouse anti synaptotagmin 2 (1:1000; Developmental Studies Hybridoma Bank ZNP-1); mouse anti-synaptotagmin 10 (1:1000; Absolute Antibodies Ab02161-1.1 clone [N268/73]); and rabbit anti-IGF-1R (1:500; Abcam AB182408).

### IESS Patients: Immunohistochemistry

Analysis of brain tissue resected during epilepsy surgery was approved by Baylor College of Medicine’s IRB. Signed informed consent was obtained from parents/legal guardians. Four patients with a diagnosis of IESS underwent epilepsy surgery at Texas Children’s Hospital. We compared IESS immunohistochemical findings to those of four “control” patients who had brain tumors and associated focal seizures and underwent tumor resections (see Supplemental Table 1 for patient details). Once resected, tissue was fixed and imbedded in paraffin.

Immunohistochemistry for human tissue (10 µm thick sections) was carried out as in rodents, but with added steps to dewax and rehydrate paraffin sections and antigen retrieval. Our synaptotagmin 2 antibody did not cross-react with the human specimens and thus we do not report results for this molecule.

### Western Blotting

The somatosensory cortex was dissected at 4°C. To accomplish this, a series of one-millimeter coronal slices (Zivic Instrument BSMAA001-1) was obtained from the cortex. Under a stereomicroscope and progressing caudally, the somatosensory cortex was identified in slices containing the first appearance of the hippocampus and 2 mm punches (Fine Science Tools 20830-00) were obtained. Tissue was immediately frozen on dry ice. Afterwards, the tissue was sonicated in a homogenizing buffer. Protein concentrations were determined using the Bradford assay (Bio-Rad, Hercules, CA) and 20μg of total protein electrophoretically separated on 4–12% gradient SDS-PAGE gels (Life-Technologies) and transferred onto PVDF membranes (Immuno-Blot PVDVF). Membranes were probed with anti-GAPDH (Millipore, CB1001-500; 1:10,000); rabbit-anti Thr308-phosphorylated-AKT (Cell Signaling, 2965S; 1:1,000); and rabbit anti-pan-AKT (Cell Signaling, 4691; 1:1,000). Near-Infrared western blot detection was performed with the Odyssey CLx Imaging system (LI-COR Biotechnology). Rabbit primary antibodies were labeled with IR Dye 800 CW, and mouse primary antibodies labeled with IR Dye 680 RD. The expression level of each protein was determined using NIH ImageJ software. Densitometry quantification was performed by normalizing the expression level of each protein to GAPDH. Levels of phosphorylated AKT were normalized to total AKT. Generated ratios were normalized to average value of the control samples on each blot and expressed as a percent of control.

### Magnetic Resonance Imaging (MRI) and cortical volume quantification

Following fixation, brains were imaged using a 9.4 T, Bruker Avance Biospec spectrometer in a 21 cm bore (Bruker Biospin, Billerica, MA) with a 35mm volume resonator. A 2D RARE sequence was used for imaging with Repetition Time (TR): 2500 ms; Echo Time (TE): 42 ms; RARE Factor: 4; NA: 2; matrix 256×256; slice thickness, 500 μm; number of slices: 23. MR images were viewed and analyzed using OsiriX DICOM viewer. To determine the volume of the cortex, a polygon tool was used to trace the perimeter of the cortex and calculate the area of the cortical hemispheres for each slice. The sum of the measured areas (∑A_i_) in combination with the inter-slice distance (750µm) was used to calculate the total volume (V) of each cortical hemisphere. Thus, for each of the brain hemispheres analyzed: V= ∑ A_i_ (0.750).

### Statistical Methods

Data are summarized as mean ± SEM. Either a one-way or two-way ANOVA was performed to determine statistical significance. A post-hoc Tukey test was used to correct for multiple comparisons or repeated measures when appropriate. When two groups were compared, a two-sample t-test was employed. For categorical data, the Fisher exact test was used. GraphPad Prism 9 (version 9.1.2) or Origin (version 8 Pro) was used to perform statistical tests.

## Results

Images in Figures 1A-C illustrate at increasing magnifications the distribution of IGF-1 in a coronal section through the somatosensory cortex (area S1 is outlined in A). We focused our analyses on area S1 since previous studies of the TTX model implicated it in spasm generation (32, 45). Three-dimensional reconstruction of immunolabeling (Figure 1D; compare to 1C) permitted quantification of the volume of cortex occupied by immunopositive cells. Our previous study of the neocortex showed that IGF-1 is colocalized with the neuronal protein MAP2 (32). However, whether IGF-1 is expressed by all neocortical neuronal subpopulations has not been addressed. To examine IGF-1 expression in GABAergic interneurons, we co-labeled sections for the GABA synthetic enzyme, Glutamic Acid Decarboxylase (GAD), and IGF-1. Images in Figures 1E-G show GAD is expressed in cells bodies (see arrows in G) and in networks of putative GABAergic nerve terminals in neocortical layers II, III IV and Vb. (For larger field of view, see Supplemental Figure 1.) Figure 1G compares GAD and IGF-1 expressions. While the distribution of the two molecules differ, merged images show GAD-positive cells co-express IGF-1. To quantify this co-expression, we isolated GAD-positive cell bodies followed by application of the Imaris colocalization algorithm (“Coloc;” see Methods). For a control (see dashed line inset), we processed histological sections without the IGF-1 primary antibody. Across animals, 46.9 ± 6.0% of GAD-immunopositive voxels co-labeled with IGF-1, compared to 0.32 ± 0.16 in control sections (Figure 1G graph).

**Figure 1.**
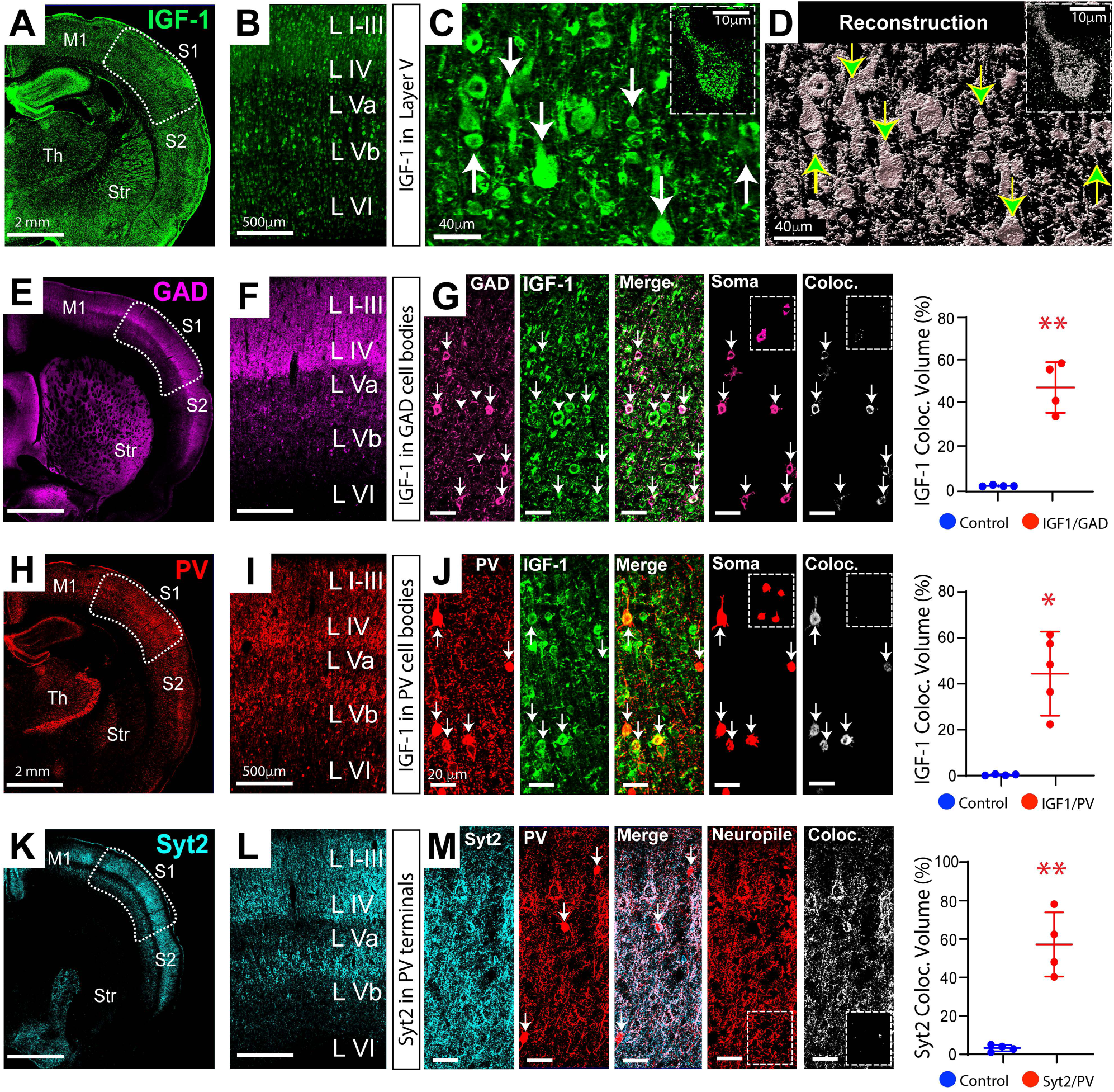
Immunohistochemical expression and distribution of IGF-1 in neocortical interneurons. (A) IGF-1 expression in a coronal section from rat neocortex; somatosensory area S1 outlined. (B and C) Area S1 shown at increasing magnifications. Inset in C shows IGF-1-containing somatodendritic vesicles in one neuron. (D) To quantify volume of tissue occupied by immunolabeling, Imaris surface-rendering and reconstruction algorithms were used to generate 3D reconstructions of immunopositive cell bodies and neuronal processes; here shown as monochrome - compare to C. Arrows denote the same cell bodies in the two panels. (E) GAD immunolabeling. (F and G left panel) Higher magnification images show GAD positive neuronal cell bodies (e.g., arrows) and networks of putative GABAergic nerve terminals (e.g., basket-like terminals at arrow heads). (G) Co-labeling of GAD neurons with IGF-1; see merged image. To quantify colocalization, GAD cell bodies (Soma) were isolated using the masking tool in Imaris and IGF-1 co-expression quantified using the Imaris “Coloc” algorithm. Plot is the percentage of GAD positive voxels that were IGF-1-positive compared to control brain sections incubated without IGF-1 antibody. Inset (dashed lines) in right-most images show sparse IGF-1 immunolabeling for control sections, reflecting background fluorescence. (H-J) PV interneurons also express IGF-1. Analysis identical to that for GAD. (K) Synaptotagmin 2 immunolabeling in neocortex. (L and M left panel) Networks of putative GABAergic nerve terminals at higher magnifications. (M) Synaptotagmin 2 co-labels PV nerve terminals; see merged image. To quantify co-localization, PV nerve terminals were isolated using masking tool (notice PV cell bodies at arrows eliminated in panel labeled neuropile) followed by “Coloc” program. Sections incubated without synaptotagmin 2 antibody served as a control; see dashed line insets in right-most images. n = 4 animals 2 males, 2 females. * p ≤ 0.05, ** p ≤ 0.01 – two sample t-test with Welsh correction. M1 - Motor Cortex, S1 and S2 - Somatosensory Cortex, Str – Striatum, Th – thalmus.

Parvalbumin (PV)-containing interneurons are a major subclass of fast-spiking neocortical neurons that play a key role in regulating network excitability (46). To augment our analysis of IGF-1 expression by interneurons, we performed the same analysis for PV cells (Figures 1H-J). Results on Figure 1J show IGF-1 is co-expressed in the cell bodies of PV interneurons.

### Subcellular Distribution of IGF-1 in Neocortical Interneurons

Little is known about the subcellular distribution of IGF-1 in interneurons, but this information should have significant functional implications. Numerous studies of neuropeptides, including IGF-1, have shown that they are localized to, and released extracellularly from, neurosecretory vesicles in cell bodies and dendrites (47, 48). While our results indicate that neocortical neurons similarly express IGF-1 in vesicles within somatodendritic compartments (e.g., insets in Figures 1C and D), it remained unclear if this growth factor is transported to presynaptic terminals for release. To explore this possibility, we pursued synaptotagmin 2 as a PV cell presynaptic marker since it has been reported to be expressed exclusively in PV nerve terminals in the visual cortex (49) and it mediates GABA release at basket cell synapses (50). In Figure 1K-M, double labeling for synaptotagmin 2 and PV showed strong co-localization in putative presynaptic nerve terminals.

Using synaptotagmin 2 as a PV cell presynaptic marker, we next co-labeled brain sections for IGF-1 but failed to observe co-localization (Figures 2A and B), indicating that IGF-1 is not transported to nerve terminals. However, in keeping with results from olfactory bulb neurons (48), we did discover that IGF-1 is localized to vesicles in the cell bodies and dendrites of interneurons and is co-expressed with vesicular synaptotagmin 10 (Figure 2C). Moreover, results showed this pairing of IGF-1 and synaptotagmin 10 was enriched in interneurons (Figures 2C-E). These findings indicate that interneurons are likely able to release IGF-1 extracellularly from somatodendritic vesicles and, as in other organs (51), this growth factor may have both autocrine and paracrine roles in neocortical development and function.

**Figure 2.**
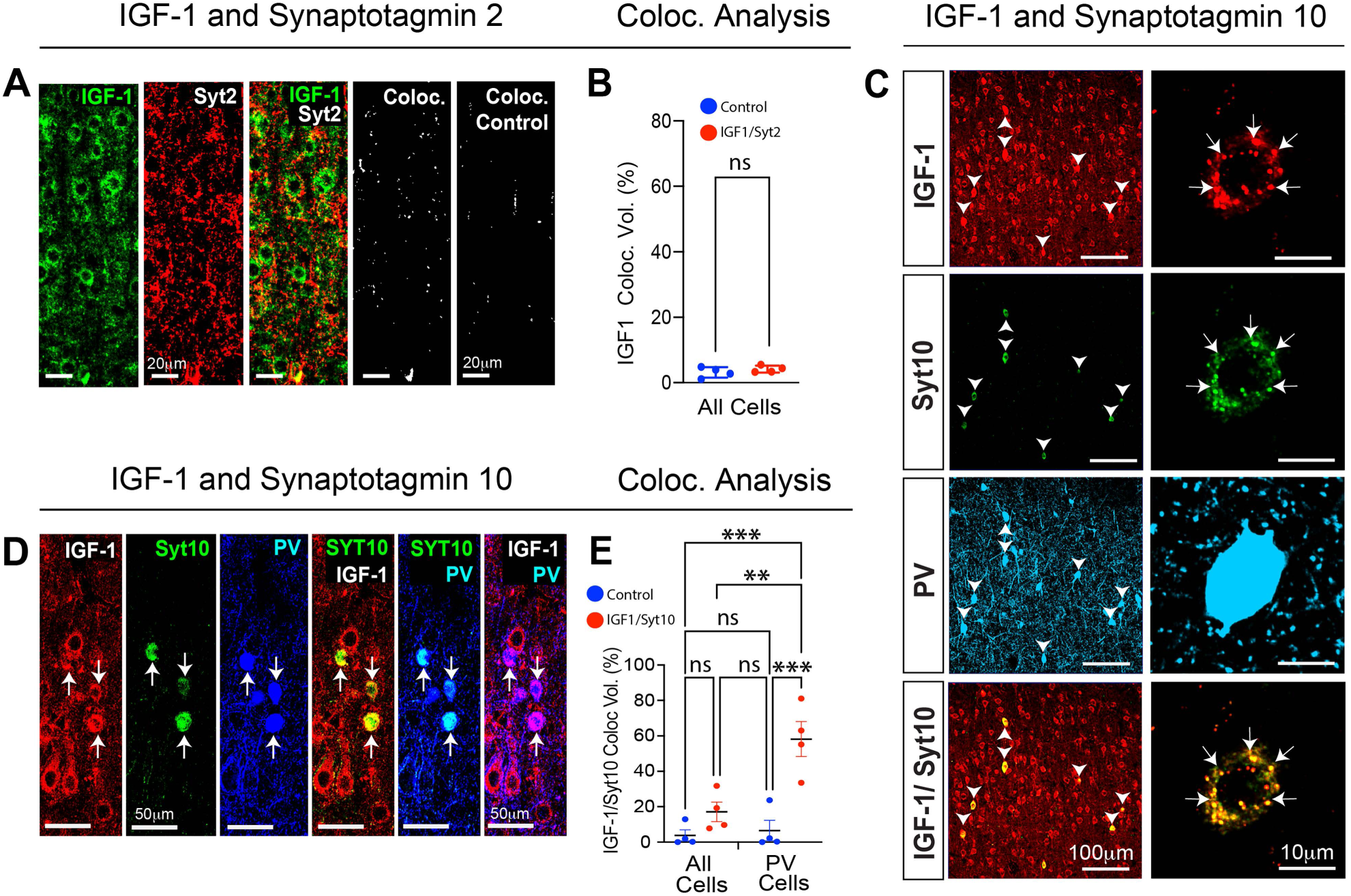
Subcellular Localization of IGF-1 in Neocortical Interneurons. (A and B) IGF-1 does not colocalize with synaptotagmin 2; see merged images and results from “Coloc” algorithm in B. Coloc Control – section incubated without the IGF-1 primary antibody reflecting background fluorescence. n = 4 animals 2 males, 2 females. ns – not significant: two sample t-test with Welsh correction. However, (C) IGF-1 and synaptotagmin 10 do co-label PV interneurons (arrow heads) and vesicles (arrows) in their somatic compartments. Moreover, (D) the pairing of IGF-1 and synaptotagmin 10 expressions is enriched in PV interneurons. Triple labeled immunohistochemistry revealed that while IGF-1 is widely expressed across neocortical neuronal populations, the expression of synaptotagmin 10 is confined to PV interneurons (see merged image Syt10/PV). This results in the enrichment of IGF-1 and synaptotagmin 10 co-expressions in these cells (see merged image Syt10/IGF-1). To quantify this enrichment, we used the “Coloc” algorithm in Imaris and first computed the colocalization of the two molecules across all cells in fields of interest and then repeated the computation after having isolated PV cells with an Imaris masking tool. (E) Quantified results indicate that for all cells the paired co-expression of IGF-1 with synaptotagmin 10 was slightly increased over controls. However, for PV cells alone co-expression was significantly enhanced over that for all cells. All Cells: 17.1 ± 5.48% vs PV Cells 58.2 ± 9.83%, p = 0.007 – two-way ANOVA with correction for multiple comparisons. ** p ≤ 0.01, *** p ≤ 0.001, n = 4 animals 2 males, 2 females. All images taken for neocortical layer V.

### Diminished Interneuron Connectivity in Epileptic Neocortex

Previously, we reported widespread loss of IGF-1 in the neocortex of the TTX model (32). Since interneurons play an important role in regulating cortical excitability and express IGF-1, we wondered if the loss of IGF-1 could impact interneurons in epileptic animals. In these experiments, TTX was chronically infused into the cortex beginning on postnatal day 12, resulting in epileptic spasms one week later that persisted for at least three months (see experimental timeline and EEG electrode placement in Figure 3A). Figure 3B compares NREM sleep EEG recordings from a TTX-infused rat and a saline-infused litter mate control, illustrating the differences in cortical excitability. Figure 3C’s raster plot illustrates the clustering of spasms over 24 hours and Figure 3D’s bar graph quantifies average daily spasm counts.

**Figure 3.**
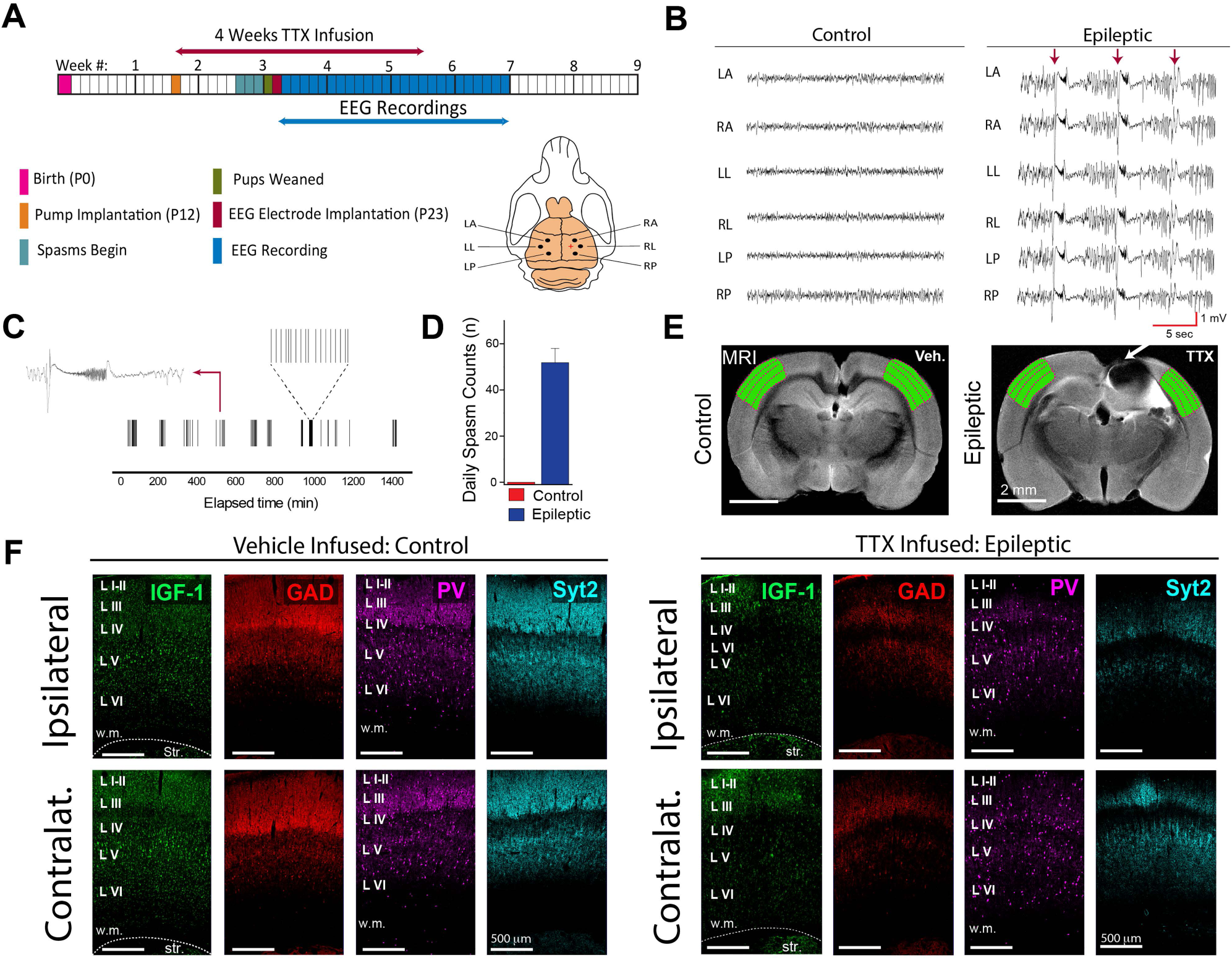
The expression of IGF-1 and GABAergic neuronal and synaptic proteins are reduced in the neocortex of animals with epileptic spasms. (A) Experimental timeline: neocortical TTX infusion began on postnatal day (P)12 and EEG electrodes implanted on P23. Recordings began the next day and were continuous for 4-5 weeks. Electrode placement was 2 mm from TTX infusion site (red +). Locations LA - left anterior, LL - left lateral, LP – left posterior. Locations in the right hemisphere are similarly named. (B) NREM sleep EEG recordings from a TTX-infused epileptic rat and its saline-infused control. Three epileptic spasms, denoted by arrows, interrupt on-going interictal slow waves that are visibly larger than those in controls. (C) Spasm clustering over 24 hrs in one rat: left inset - an individual ictal event; right inset - occurrence of individual spasms in one cluster. (D) Epileptic animals had 51.8 ± 6.2 daily spasms over 5 days (P40-45), controls had none. (E) MRI images from saline and TTX-infused rats (TTX-induced lesion demarcated by arrow); the adjacent area S1 and contralateral counterpart are shaded green as is S1 in the control animal. (F) Immunohistochemical examination of area S1 both ipsilateral and contralateral to TTX infusion shows diminished expression of the 4 molecules studied in epileptic animals compared to the control. n = 6 for both groups: 3 males, 3 females. Syt2 - synaptotagmin 2, Str - striatum

In the TTX model, an apoptotic-mediated lesion is formed at the TTX infusion site (32). MRI images in Figure 3E depict the lesion (see arrow) and its relationship to the adjacent somatosensory area S1, shaded green. Figure 3F compares levels of IGF-1, GAD, PV, and synaptotagmin 2 in area S1, both ipsilateral and contralateral to the TTX lesion to their counterparts in saline-infused rats. Confirming our previous results, IGF-1 levels were reduced in both regions of epileptic rats. Importantly, in these same areas we observed a decrease in the expression of PV, GAD, and synaptotagmin 2. Higher magnification images in Figures 4A and B illustrate expression at the cellular level and a reduction in the density of putative inhibitory interneuron nerve terminal networks. When quantified, immunolabeling was reduced (Figures 4C and D: see also Supplemental Figure 2 for representative 3D reconstructions used to quantify the differences in immunoreactivity). However, the number of interneuron cell bodies was unchanged (Figures 4D and E).

**Figure 4.**
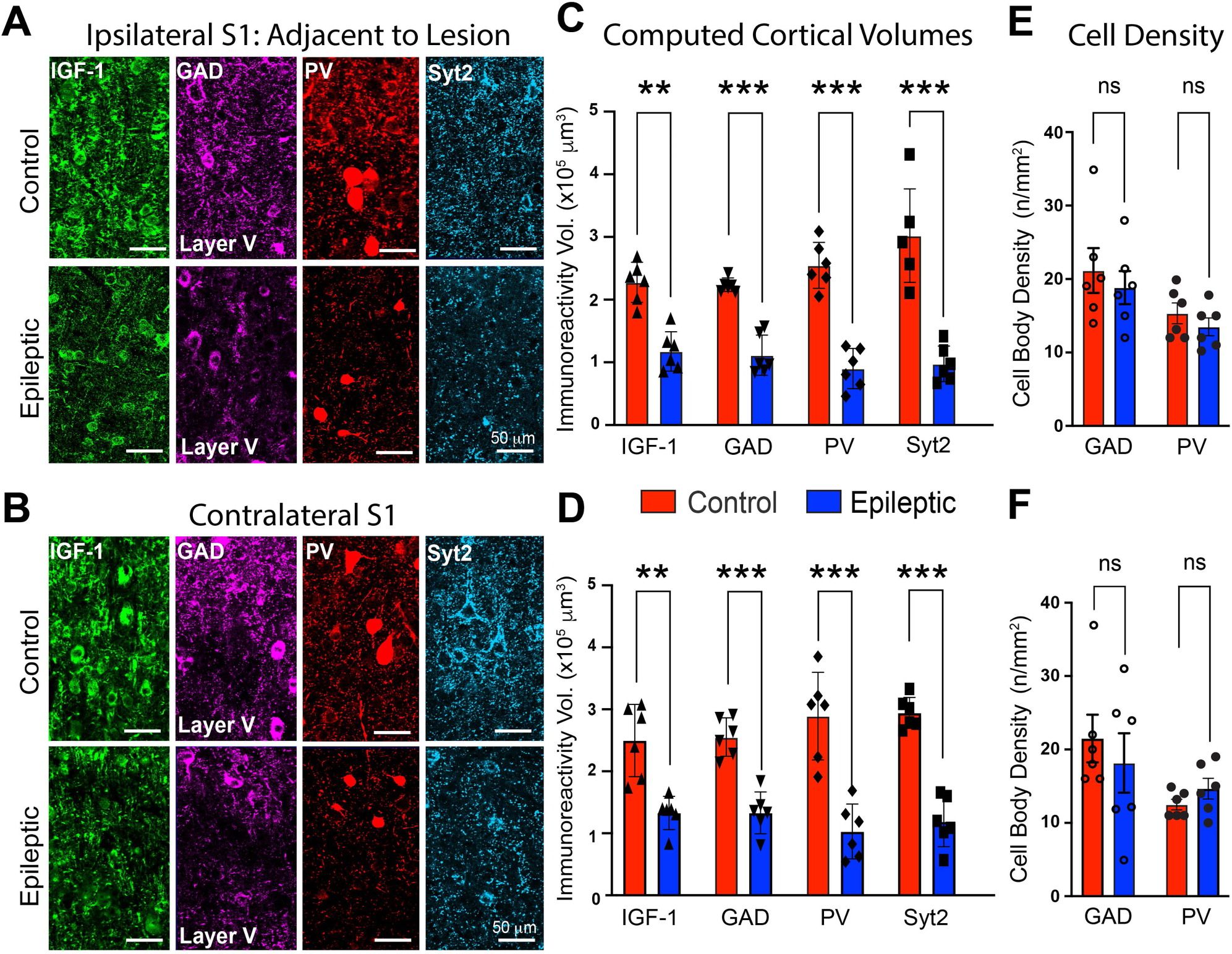
Analysis of IGF-1, GAD, PV, and synaptotagmin 2 in areas S1. (A and B) Higher magnification images of area S1 in epileptic rats show a decrease in proteins expressed by putative GABAergic nerve terminals yet the number of immunolabeled interneurons is unchanged. IGF-1 expression is also suppressed in the epileptic neocortex both ipsilateral and contralateral to TTX-induced lesion. Note the relatively sparce putative presynaptic nerve terminal labeling by all three GABAergic proteins in the epileptic neocortex. (C and D) Volumetric quantification of immunoreactivity performed with Imaris surface rendering/reconstruction algorithm. (E and F) Neuron cell body density computed with Neurolucida. N=6 for both groups: 3 males, 3 females. ** p ≤ 0.01, *** p ≤ 0.001, ns non-significant - two-way ANOVA corrected for multiple comparisons.

### Loss of Neocortical IGF-1 is Largely Limited to GAD-Containing Interneurons

Examination of the doubled labeled images in Figures 1G and J shows that IGF-1 is co-expressed in interneurons with either GAD or PV. However, other cells in these images are clearly IGF-1 positive. Since our previous studies (32) have shown that IGF-1 co-labels essentially all cells with the broad neuronal marker MAP2, we surmised that the majority of the IGF-1-positive, GAD and PV-negative cells are neocortical pyramidal cells. Given that IGF-1 expression is suppressed in the epileptic neocortex we next wondered if its expression was preferentially diminished in inhibitory or excitatory neurons. To address this question, we co-labeled neocortical sections with IGF-1 and GAD. Using the Imaris masking tool to identify cells expressing GAD, we were able to quantify IGF-1 expression in these interneurons. We also quantified IGF-1 in the entire field of view and by subtracting IGF-1 expression levels in interneurons from that in all cells we were able to estimate the loss of IGF-1 from GAD-negative cells. Results in Figure 5 show that IGF-1 was preferentially lost from interneurons. The merged images in the lower panels in Figure 5A indicate that while most GAD-positive interneurons in control animals express high levels of IGF-1 (arrow heads - lower panels), interneuron IGF-1 expression in epileptic rats is either absent or in much lower expression levels. Indeed, our analysis in Figure 5B suggests that while IGF-1 levels in GAD-containing interneurons was markedly reduced, IGF-1 expression in GAD-negative cells was unaltered. These results could implicate impaired local autocrine IGF-1 signaling in interneuron dysfunction in epileptic neocortex (see Discussion).

**Figure 5.**
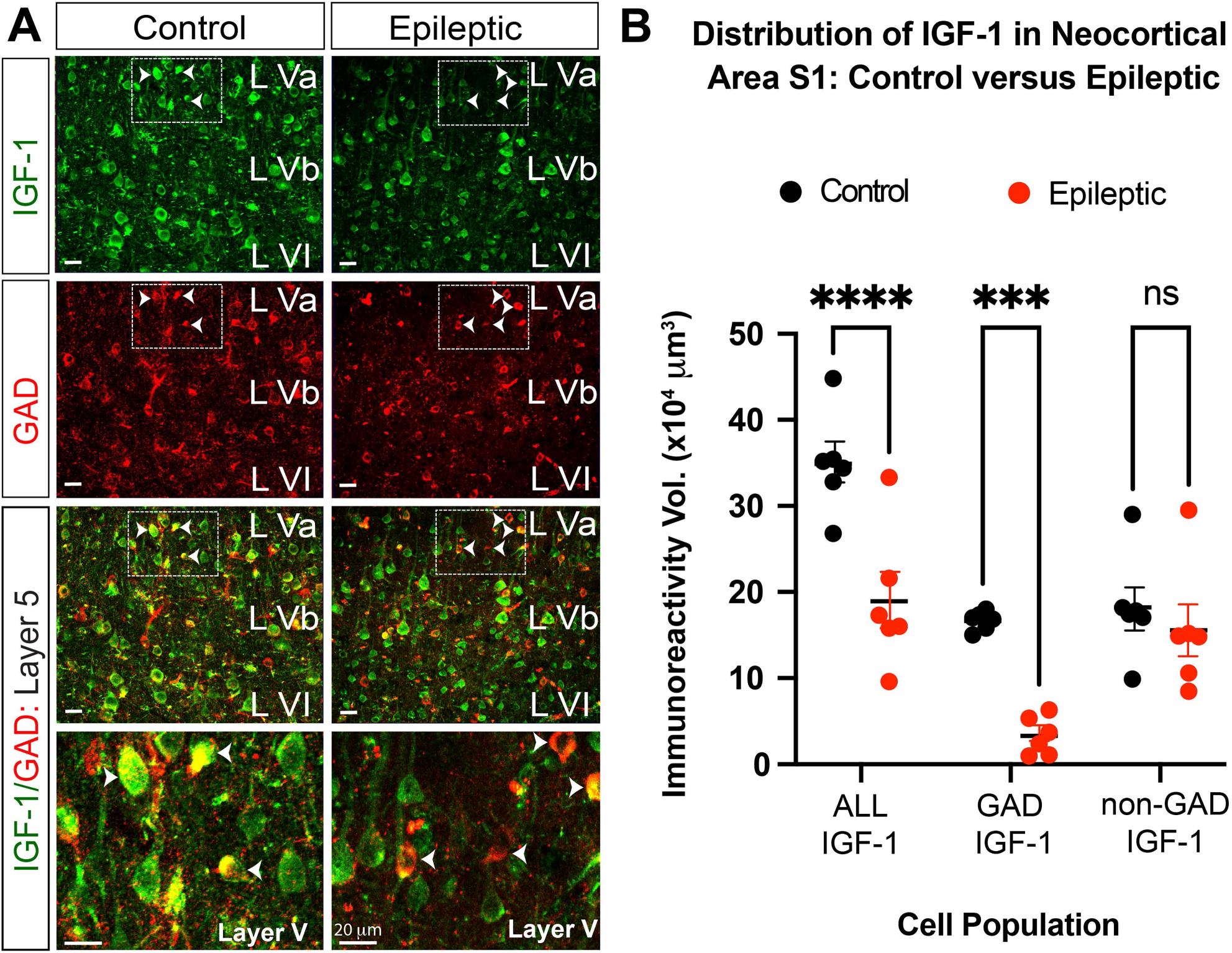
Loss of IGF-1 in epileptic neocortex is predominantly confined to GAD-containing interneurons. (A) Both GAD and IGF-1 expression are reduced in epileptic animals. However, merged images indicate that the loss of IGF-1 was not uniform across all neurons. In control rats, IGF-1 is expressed at high levels in most GAD-containing cells (yellow in merged images) but in epileptic rats IGF-1 levels in the interneurons are lower or absent. Boxed areas (dashed white lines) in lower magnification IGF-1, GAD panels and merged images are shown at higher magnification in lowest panels. Arrow heads denote the same neurons in each panel from control and epileptic animal. (B) Comparison of cortical volumes occupied by IGF-1 in all cells and GAD positive interneurons. IGF-1 expression in GAD-negative cells was computed by subtracting these two values from each other for each animal. While IGF-1 is markedly reduced in GAD-positive cells, IGF-1 in GAD-negative cells appear unchanged. Volumes were computed for area S1 in both hemispheres and averaged for each animal. Cortical volumes were computed using Imaris surface rendering/reconstruction algorithms. N = 6 for both groups: 3 males, 3 females. ** p ≤ 0.01, *** p ≤ 0.001, **** p ≤ 0.0001, ns – non-significant – two-way ANOVA corrected for multiple comparisons.

### Interneuron Connectivity Reduced in Cortex of IESS Patients

Having observed the loss of molecules expressed by inhibitory interneurons and their presynaptic nerve terminals in the TTX model, we sought to determine if similar changes take place in children with IESS. To do this, we obtained surgically resected neocortical tissue from four IESS patients who had undergone surgery to control their seizures. At surgery, two of the children continued to have epileptic spasms and the other two had focal motor seizures. All had a history of perinatal strokes. For comparison, we examined cortical tissue from four patients that was adjacent to tumors removed during cancer surgery (see Supplemental Table 1 for patient details). Compared to the tumor control tissue, we observed a dramatic decrease in the expression of GAD, PV, and IGF-1 in spasm patients, marked by a decrease in the density of putative GABAergic nerve terminals (Figures 6A-C).

**Figure 6.**
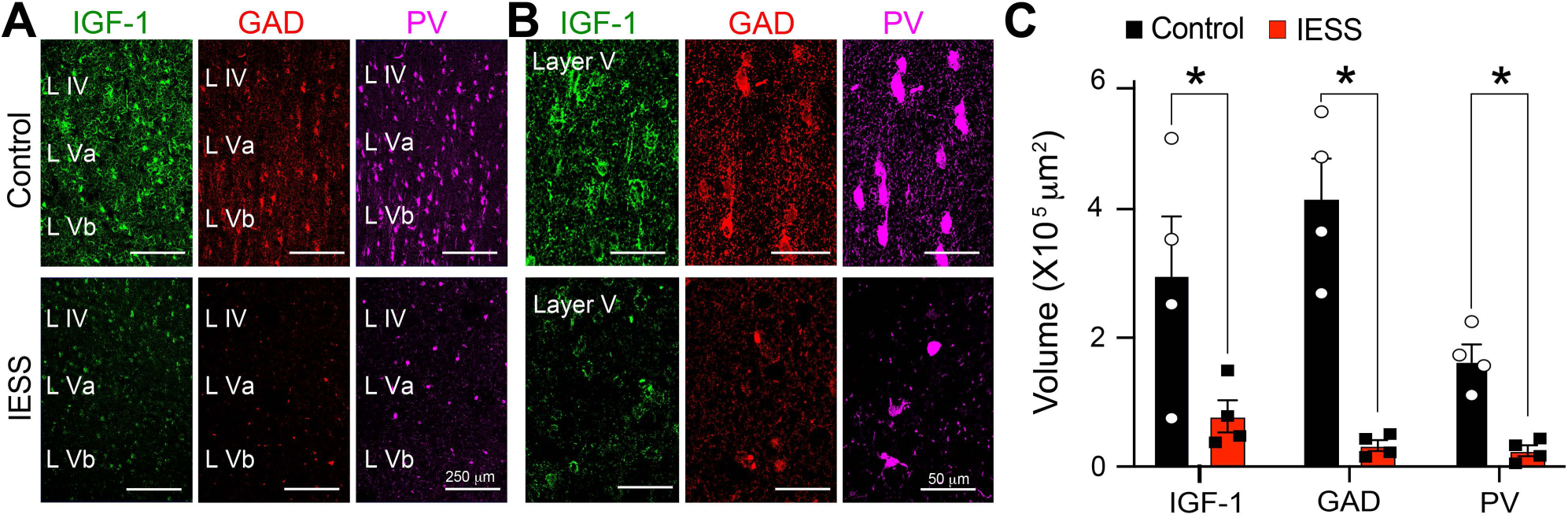
Alterations in the expression of IGF-1, GAD, and PV in the neocortex of IESS patients. (A) Compared to resected neocortical tissue adjacent to tumors in control patients, expression of GAD, PV, and IGF-1 were reduced in surgically resected tissue (adjacent to frank GFAP-positive gliotic neocortical lesions) in IESS patients. Interneuron cell bodies appear noticeably smaller in epileptic tissue. (B) At higher magnification, sparse labeling of putative GABAergic nerve terminals is apparent in sections from spasm patients. (C) Quantification of results with Imaris – surface rendering and reconstruction algorithms. * p ≤ 0.05 - two-way ANOVA corrected for multiple comparisons.

### Knock-down of IGF-1R in Neonatal Neocortex Reduces Interneuron Connectivity

Brain-derived IGF-1 has well-known neuronal growth-promoting properties in the brain (25). Thus, we wondered if our observed reduction in IGF-1 expression in the cortex of epileptic rats could impact interneuron development and contribute to the observed decreases in interneuron synaptic proteins and concomitant diminished connectivity. PV expression is known to increase dramatically during the first few postnatal weeks in rodents (52). While dramatic developmental changes have also been observed in GAD expression (53), developmental studies of synaptotagmin 2 in somatosensory cortex have not been reported. Figures 7A and B show that synaptotagmin 2 expression closely parallels changes in PV. Both molecules are very low at birth but show marked increases in expression over the next three to four weeks – featured by a notable increase in the density of nerve terminal networks across the neocortical laminae.

**Figure 7.**
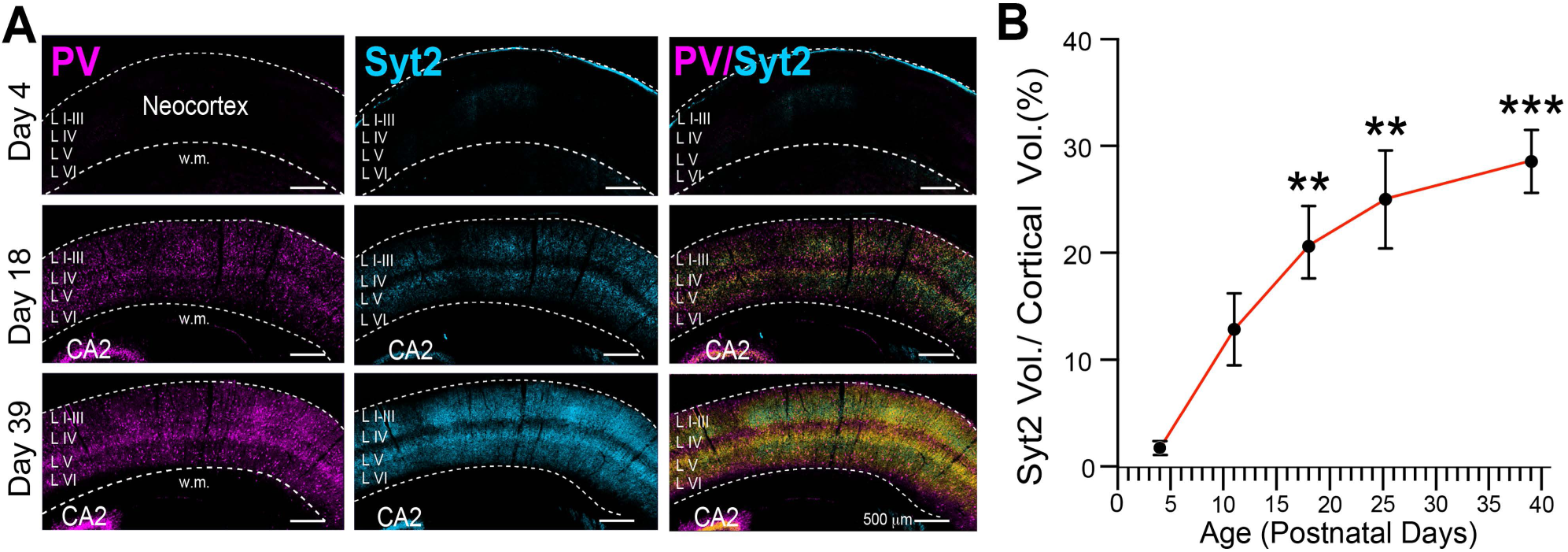
Synaptotagmin 2 immunohistochemistry in the mouse neocortex reveals the postnatal formation of GABAergic nerve terminal networks. (A) Postnatal increases in synaptotagmin 2 immunolabeling closely parallel that of PV in the mouse somatosensory cortex and reveals the dramatic developmental increase in putative GABAergic presynaptic nerve terminal networks arising from PV neurons. (B) Summary of developmental increases in synaptotagmin 2 immunolabeling during the first 5.5 weeks of postnatal life. To account for neocortical growth, immunolabeled synaptotagmin 2 volumes are expressed as a percent of tissue volume analyzed. Statistical comparisons were made to synaptotagmin 2 expression on P4. ** p ≤ 0.01, *** p ≤ 0.001 - one-way ANOVA corrected for multiple comparisons. CA2 - hippocampal subfield, w.m. - white matter.

To address the hypothesis that a reduction of IGF-1 in the epileptic neocortex impairs the development of inhibitory interneurons, we sought to eliminate the receptor for IGF-1 (IGF-1R) from neocortical neurons soon after birth. Prior studies suggested that IGF-1R is expressed by all neurons of the brain. However, since inhibitory interneurons are a minority of neurons in the CNS, we first sought to verify that they express IGF-1R. Images in Figure 8A show that essentially all PV- and GAD-containing neurons also co-express IGF-1R. With this information, we set out to knock-down IGF-1R expression in conditional IGF-1R^lox^ knockout mice by a unilateral focal injection of AAV-Cre virus into the neonatal neocortex (Figures 8B and C). IGF-1R^lox^ mice injected with control AAV lacking Cre and naïve mice served as controls. Figures 8D and E show that by P30 IGF-1R expression is greatly reduced in neocortical regions expressing the AAV-Cre reporter compared to both controls. Importantly, we also observed a reduction in the GAD, PV, and synaptotagmin 2 synaptic networks that had developed by this age. Thus, a reduction in IGF-1 signaling through its receptor appears to impact the maturation of inhibitory interneurons and GABAergic synaptogenesis. This led us to suspect that the loss of IGF-1 in the epileptic cortex impaired interneuron development and this contributed at least in part to the loss of neocortical inhibitory interneuron connectivity.

**Figure 8.**
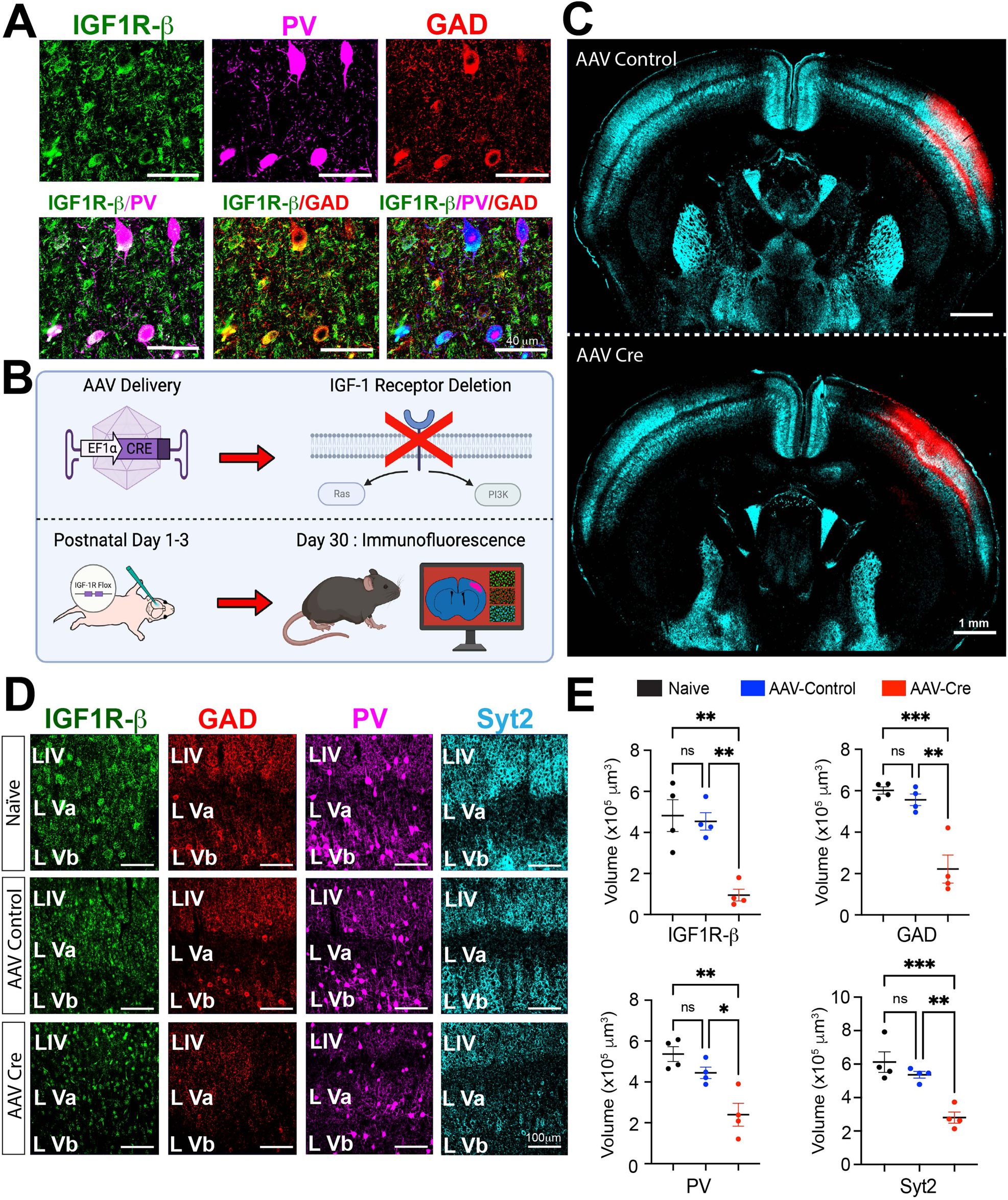
Neocortical GABAergic interneurons express the receptor for IGF-1 and its elimination neonatally leads to a decrease in neocortical expression of GABAergic synaptic proteins one month later. (A) Immunohistochemistry for IGF-1R, GAD, and PV in the mouse neocortex showing colocalization of IGF-1R with both interneuron proteins in double- and triple-labeled merged images. (B) Schematic of experimental design: a focal unilateral intracortical injection on P1, 2 or 3 of AAV-Cre or AAV control virus without Cre in IGF-1R^lox^ mice, followed by immunohistochemistry on P30. (C) Images showing expression of the AAV reporter for the AAV-Cre and AAV-Control viruses at neocortical injection sites. Synaptotagmin 2 served as a counterstain. (D) Comparison of IGF-1R, GAD, PV, and Synaptotagmin 2 (SYT2) expressions in the neocortex of naïve mice (not injected with virus) and neocortical regions transfected with AAV-Cre or AAV-Control without Cre. (E) Volumetric quantification of results across animals with Imaris – surface rendering and reconstruction algorithms. N = 4 per group 2 males, 2 females but for AAV-control 3 males and 1 female * p ≤ 0.05, ** p ≤ 0.01, *** p ≤ 0.001, ns non-significant - one-way ANOVA corrected for multiple comparisons.

### Treatment with (1–3)IGF-1 Abolishes Spasms and Rescues Interneuron Connectivity

Since decreases in neocortical GABAergic synaptic networks could contribute to the generation of epileptic spasms, we reasoned that experimental strategies that would restore IGF-1 levels might also renew interneuron development, rescue GABAergic synaptic connectivity, and possibly abolish spasms. One way to test this hypothesis would be to treat epileptic animals with IGF-1. While IGF-1 has been reported to cross the blood-brain barrier (54), its lack of success in clinical trials for neurological disorders has been disappointing. Only dosages below 0.25 mg/kg/day are approved by the FDA due to concerns over side effects. We performed a dose response experiment for rhIGF-1 in mice and used phosphorylation of AKT in the neocortex as a readout of its effects. Dosages at and above 1 mg/kg activated the IGF-1R downstream effector AKT. However, the clinically approved dosage of 0.25 mg/kg was unable to activate AKT (Figure 9A), which could help explain its clinical ineffectiveness and on spasms in the TTX model at this dosage (32).

**Figure 9.**
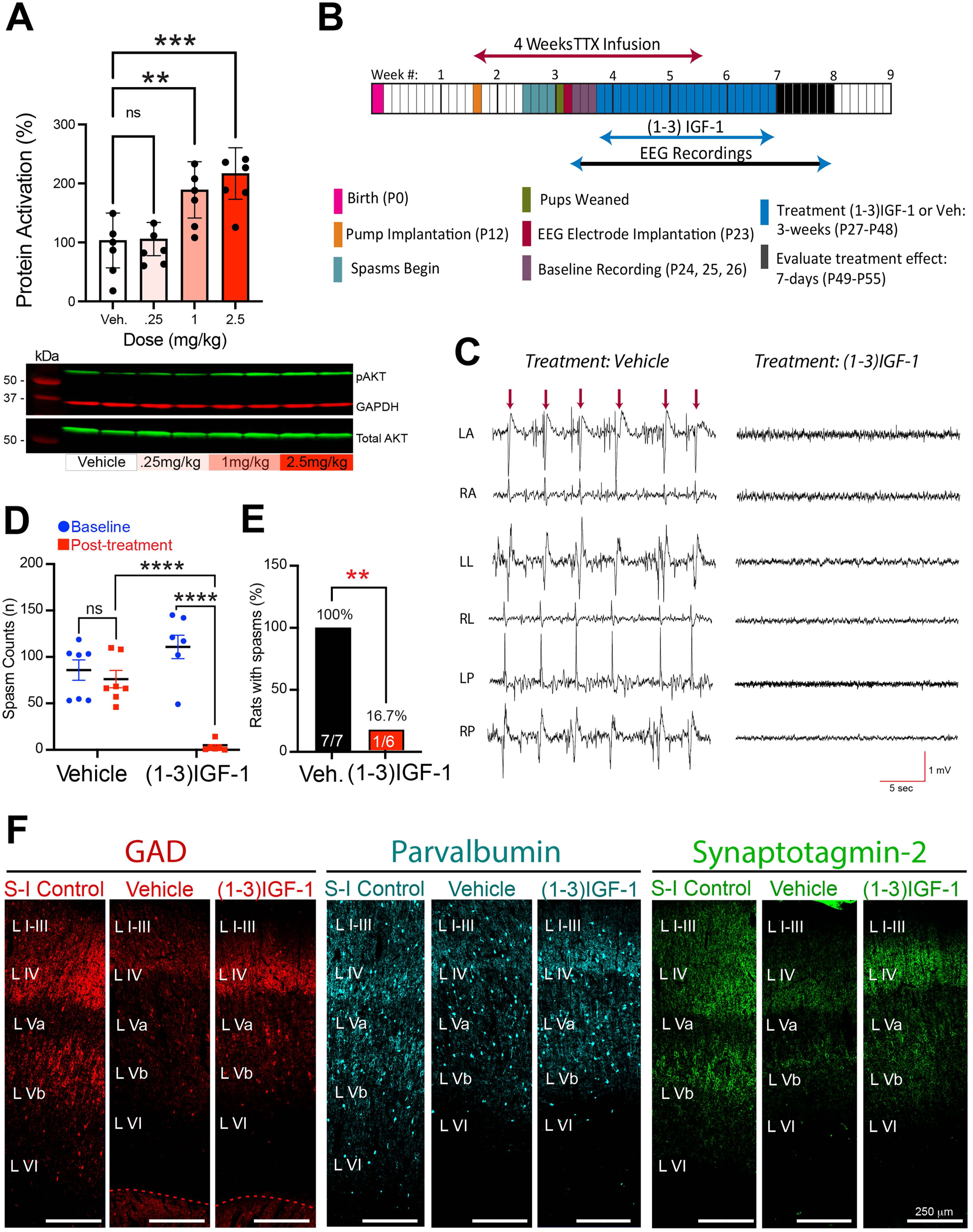
Resolution of epileptic spasms and rescue of GABAergic synaptic proteins after treatment with the IGF-1 derived tripeptide (1–3)IGF-1. (A) rhIGF-1 fails to activate the neocortical IGF-1R downstream effector AKT in the mouse neocortex at the FDA-approved dosage (0.25 mg/kg). Responses to single i.p. injections. Cortex collected 2 hrs later. (B) Experimental timeline and (1–3)IGF-1 treatment protocol: neocortical TTX infusion began on P12, EEG electrodes implanted on P23. Recordings began on P24 and after 3 days of baseline recordings, treatment with (1–3)IGF-1 began and continued for 3 weeks. Treatment - 40 mg/kg/day by gavage, controls received vehicle. Effects on spasms evaluated the week after treatment. (C) Representative recordings following treatment: (1–3)IGF-1-treated rat was spasm-free but vehicle-treated rat continued to have frequent spasms; here, 6 spasms (arrows) within a prolonged cluster are shown. Recordings were taken during waking. (D) Summary comparing baseline and posttreatment daily spasm counts in vehicle and (1–3)IGF-1 treatment groups. Posttreatment daily spasm counts: vehicle: 74.5 ± 9.3 vs (1–3)IGF-1 3.30 ± 2.21. n=7 for vehicle, and 6 for (1–3)IGF-1. Males = 8 Females = 5. ** p ≤ 0.01, *** p ≤ 0.001, ns non-significant - two-way ANOVA corrected for multiple comparisons. (E) All vehicle-treated rats continued to have spasms posttreatment. Only one (1–3)IGF-1-treated animal had spasms. Statistical comparison - Fisher Exact Test (F) Treatment with (1–3)IGF-1 rescued GAD, PV and synaptotagmin 2 expressions. S-I – Saline-Infused Control.

As an alternate treatment strategy, we turned to (1–3)IGF-1, the n-terminal tripeptide of IGF-1. We have reported that (1–3)IGF-1 can abolish spasms in a majority of epileptic animals and provided evidence that it activates IGF-1R (32). Thus, we treated epileptic rats with (1–3)IGF-1, first assessing its effects on spasms (see Figure 9B for experimental timeline) and then examining alterations in neocortical expression of the three interneuron biomarkers. Figure 9C compares recordings from a vehicle- and a (1–3)IGF-1-treated rat in the week following treatments. While all TTX-infused rats (seven out of seven) treated with vehicle continued to have spasms, the majority of (1–3)IGF-1-treated rats (five of six) were seizure-free (Figure 9E). (1–3)IGF-1 produced a 96% decrease in daily spasm counts compared to vehicle-treated epileptic rats (Figure 9D).

When we compared immunohistochemical results in vehicle- and (1–3)IGF-1-treated rats to those from saline-infused non-epileptic controls, we found that treatment with (1–3)IGF-1 rescued GAD, PV, and synaptotagmin 2 expression to an extent comparable to non-epileptic controls. Figure 9F compares images taken from area S1 adjacent to the TTX lesion to its counterparts in the two control groups. Higher magnification images in Figures 10A and B reveal the rescue of the dense network of GABAergic nerve terminals, both ipsilateral and contralateral to the TTX-induced lesion, which is quantified in Figures 10C and D respectively.

**Figure 10.**
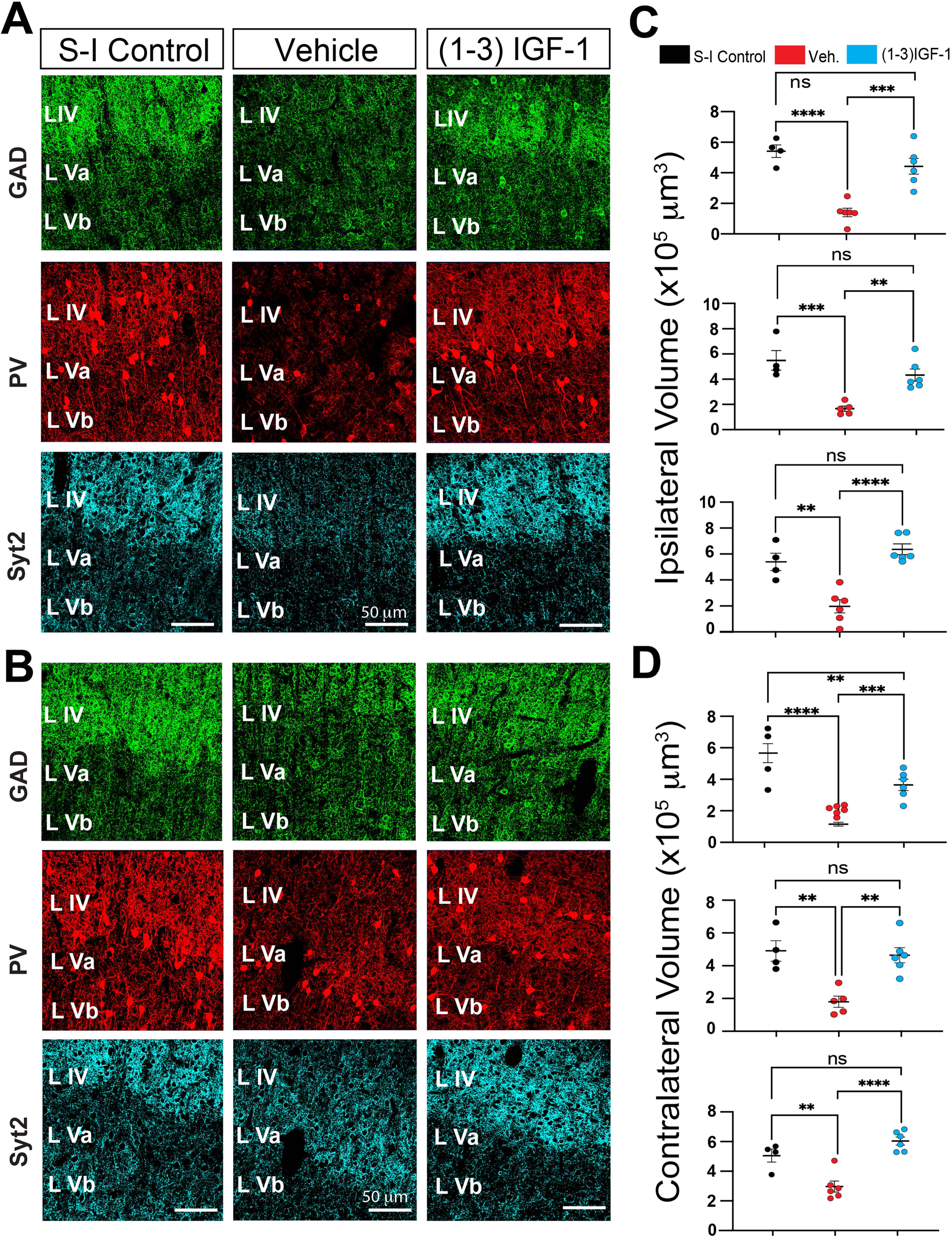
Analysis of GAD, PV, and synaptotagmin 2 expressions at higher magnification supports the conclusion that (1–3)IGF-1 treatment rescues neocortical GABAergic nerve terminal networks in epileptic rats. (A and B) Comparison of immunohistochemical results in non-epileptic saline-infused control rats, and epileptic TTX-infused rats treated with either (1–3)IGF-1 or its vehicle. Images from neocortical area S1 (ipsilateral A and contralateral B to infusion site) are shown. Note the density of putative GABAergic nerve terminals is reduced in the epileptic neocortex of animals treated with vehicle but after (1–3)IGF-1 treatment, presynaptic nerve terminal density approaches that seen in saline-infused controls. (C and D) Volumetric quantification of GAD65/67, PV, and synaptotagmin 2 immunolabeling performed by the surface rendering and reconstruction algorithms in Imaris. ** p ≤ 0.01, *** p ≤ 0.001, **** p ≤ 0.0001, ns non-significant - one-way ANOVA corrected for multiple comparisons. n = 7 for vehicle, 6 for (1–3)IGF-1 and 4 for controls. S-I – Saline-Infused Control.

One potential explanation for the rescue of inhibitory interneuron connectivity by (1–3)IGF-1 could be that the tripeptide limited TTX-induced apoptosis and smaller lesions resulted in the observed increase in interneuron connectivity. However, measurements of neocortical volumes from whole brain MRI images indicated that (1–3)IGF-1 did not alter lesion size (see Supplemental Figure 3)

### Treatment with (1–3)IGF-1 Restores Neocortical Levels of IGF-1

At this point, our results supported the hypothesis that treatment with (1–3)IGF-1 leads to activation of IGF-1R, which in turn rescues interneuron connectivity and abolishes epileptic spasms. However, it was unclear how (1–3)IGF-1 activated IGF-1R, but since IGF-1 is the agonist for IGF-1R, we examined IGF-1 expression in the epileptic rats treated with the tripeptide and their controls. Results in Figures 11A-C revealed a restoration of IGF-1 expression, both ipsilateral and contralateral to the TTX lesion. Higher magnification images in Figure 11B suggest IGF-1 expression in individual neocortical neurons is restored to a level comparable to that in saline-infused non-epileptic controls. Thus, restoration of neocortical levels of IGF-1 by (1–3)IGF-1 is likely responsible for rescuing GABAergic interneuron connectivity.

**Figure 11.**
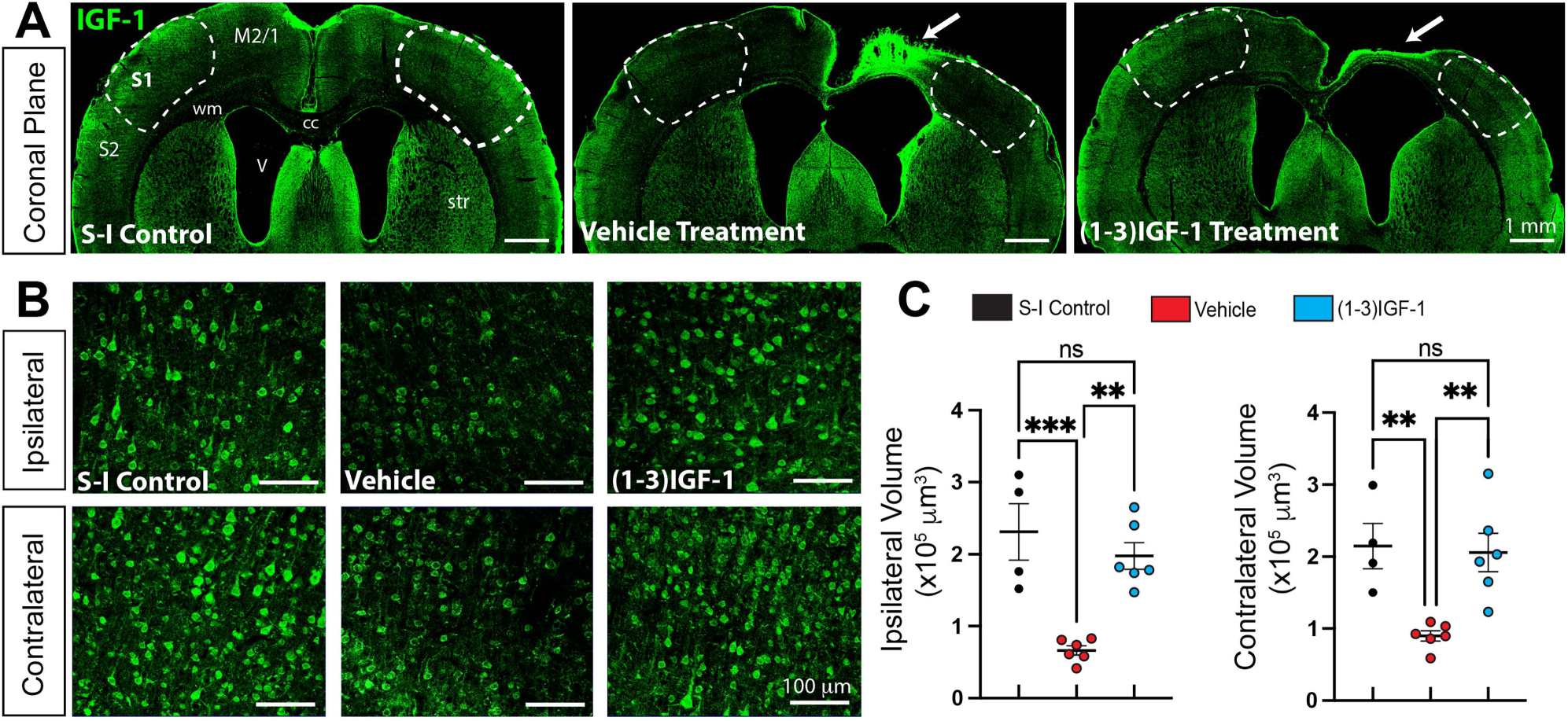
Treatment with (1–3)IGF-1 restores IGF-1 levels in the neocortex of epileptic rats. (A) IGF-1 immunolabeling in coronal sections from a non-epileptic saline-infused control rat and TTX-infused epileptic rats treated with either vehicle or (1–3)IGF-1. As reported previously, IGF-1 levels are increased within the TTX lesion (see arrows) but elsewhere in the cortex of vehicle-treated rats, including areas S1 (outlined by dashed lines), IGF-1 expression is suppressed. IGF-1 levels approach that of the saline-infused control after (1–3)IGF-1 treatment. (B) Higher magnification of IGF-1 immunolabeling of area S1. (C) Volumetric quantification of IGF-1 immunolabeling done using surface rendering and reconstruction algorithms of Imaris. ** p ≤ 0.01, *** p ≤ 0.001, ns non-significant - one-way ANOVA corrected for multiple comparisons. n = 7 for vehicle, 6 for (1–3)IGF-1 and 4 for controls. Motor Cortex -M2/1, Somatosensory Cortex – S1 and S2, wm – white matter, str – striatum, cc – corpus callosum, V ventricle, S-1 – Saline-Infused Control.

## Discussion

Our efforts focused on understanding potential relationships between neocortical IGF-1 deficiencies and interneuron abnormalities in the generation of epileptic spasms. Since our previous study (32) showed that IGF-1 expression is reduced in the neocortex of the TTX animal model and having discovered that IGF-1 is expressed by neocortical interneurons (Figure 1), we explored whether inhibitory interneurons were impacted in epileptic animals. Results in Figures 3 and 4 show that inhibitory interneuron connectivity was reduced along with the expression of the synaptic proteins. Thus, it seems likely that GABAergic inhibition is reduced in epileptic animals.

### Interneuron Dysmaturation as a Potential Mechanism Underlying Spasms

When considering possible mechanisms that could explain both injury-induced loss of interneuron connectivity and its rescue by (1–3)IGF-1 (Figure 9 and 10) there are limited possibilities. Nonetheless, one likely mechanism is interneuron dysmaturation or arrested development (55, 56). Primary neuronal dysmaturation has been observed in several brain regions, including the neocortex (57, 58), and has engendered significant interest since it has been postulated that dysmaturation might be reversed by “restorative interventions” (55, 56). In these experimental systems, brief periods of prenatal hypoxemia or hypoxia-ischemia resulted in diminished neuronal connectivity, but not neuronal death. While there are differences between the studies reported here and studies of dysmaturation following prenatal brain injury, both report decreases in synaptic connectivity after an insult to the brain early in life. An alternative explanation for the loss of GABAergic synaptic proteins and connectivity could be interneuron cell death. However, we did not observe a decrease in interneuron numbers (Figures 4E and F). Moreover, the rescue of GABAergic synaptic proteins after treatment with (1–3)IGF-1 is in keeping with dysmaturation as a mechanism but inconsistent with cell death.

Assuming that interneuron dysmaturation contributes to loss of connectivity, we next focused on the molecular events that might produce it. It is well known that brain-derived growth factors play important roles in neuron development. Since we had observed decreased neocortical expression of IGF-1 in epileptic rats, we wondered if the loss of IGF-1 might contribute to interneuron dysmaturation. With this in mind, we partially eliminated IGF-1R from neonatal cortices of IGF-1R^lox^ mice (Figures 8B-E) and by P30, we observed significant decreases in the expression of GAD, PV, and synaptotagmin 2. Since interneurons grow and form synaptic connections between P0 and 30 (Figures 7A and B), these results implicate the decrease in IGF-1 levels and resulting impairment of interneuron maturation in the loss of the interneuron presynaptic markers and connectivity in the cortex of epileptic rats.

### (1–3)IGF-1 Unexpectedly Restores Neocortical IGF-1 Expression

To further explore whether IGF-1 loss in the epileptic neocortex contributed to diminished GABAergic synaptic proteins, we considered IGF-1 replacement strategies. While treatment with rhIGF-1 was an obvious first choice, our results in Figure 9A suggested that, due to constraints of the blood-brain barrier, rhIGF-1 is only able to activate the IGF-1 signaling pathway in the neocortex at dosages reported to produce significant peripheral side effects such as hypoglycemia (59). Thus, we turned to (1–3)IGF-1, which has been reported to have neurotropic properties similar to IGF-1 (60) and to cross the blood-brain barrier (61). In an earlier study, we provided three lines of evidence showing that when (1–3)IGF-1 increases phosphorylation of AKT in the brain, it acts through the IGF-1R (32). These included results showing that Cre-virus-induced reduction of the expression of IGF-1R from the neocortex of conditional IGF-1R knock-out mice reduced activation of the IGF-1R downstream effector AKT and pretreatment with the IGF-IR antagonist, PPP, similarly suppressed (1–3)IGF-1 phosphorylation of AKT. In addition, treatment with (1–3)IGF-1 led to phosphorylation of the β-subunit of IGF-1R itself. However, at that time it was unclear how this was accomplished since it was very unlikely that (1–3)IGF-1 would bind to IGF-1R. Results in Figure 11 provide an explanation since treatment with this tripeptide restored IGF-1 levels in the cortex of epileptic rats.

### Potential Roles for Interneuron IGF-1 Autocrine Signaling in Epileptic Spasm Generation and Resolution

The molecular mechanism through which (1–3)IGF-1 increases IGF-1 expression is unknown. While little is known about IGF-1 metabolism in the brain, more is known about IGF-1 in the periphery. Circulating IGF-1 is synthesized and released from the liver to mediate somatic growth (51). However, it is also synthesized in other organs, where it has local autocrine and paracrine roles (51, 62). These mechanisms may be particularly important in the brain since IGF-1 entrance from the blood (54) is restricted by the blood-brain barrier. There is good evidence that brain-derived IGF-1 plays a role in brain growth (25–28). Studies have shown that IGF-1 is synthesized by CNS neurons (63), stored in neurosecretory vesicles and released extracellularly by synaptotagmin 10 (48). Immunohistochemical studies in Figures 2C-E are consistent with these findings. Thus, it is conceivable that IGF-1 has unappreciated autocrine and paracrine roles in neocortical development. With this in mind and given our results that IGF-1 expression in neocortex is largely restricted to GAD-positive interneurons (Figure 5), it seems plausible that impaired local IGF-1 autocrine signaling plays a role in interneuron dysmaturation in epileptic neocortex and that interneurons are consequently uniquely vulnerable to the impact of early-life brain injury. Indeed, a recent study reported that focal activation of hippocampal dendrites (by glutamate uncaging) leads to the extracellular release of IGF-1 that diffuses only a few microns from the release site (64). Such focality is perhaps not surprising since extracellular levels of IGF-1 is highly regulated not only by numerous extracellular peptidases that degrade it but also by high affinity IGF-1 binding proteins that sequester it and limit its access to IGF-1R (65). Thus, the dramatic loss of IGF-1 from interneurons would be expected to markedly reduce their ability to mediate their own development and result in their dysmaturation. On the other hand, restoring IGF-1 levels by treating with (1–3)IGF-1 could reinitiate interneuron growth and abolish spasms.

While our studies have focused on IGF-1’s role in interneuron development and function, we cannot rule out an impact of neocortical injury on glutamatergic pyramidal cells via similar mechanisms. However, since the loss of IGF-1 is largely restricted to interneurons it would appear likely that autocrine signaling in pyramidal cells in epileptic cortex is not compromised. Nonetheless, impaired local paracrine signaling from interneurons to nearby pyramidal cells would be possible. In this regard, a previous study of a Rett syndrome mouse model reported that treatments with (1–3)IGF-1 partially rescued dendritic spine density, excitatory synaptic potential amplitude and PSD95 expression (66). So, it is possible that glutamatergic synaptic transmission could be similarly impacted by treatment in the TTX model. Additional studies will be required to unravel the impact of early-life brain injury on neocortical pyramidal cell development

### Hypothetical Schemes for Spasm Generation and Resolution

In light of our findings, we propose the following schemes for the induction of epileptic spasms and their resolution by (1–3)IGF-1 (Figure 12). For induction, we posit that an injury leads to a reduction in IGF-1 levels in the neocortex which occurs prominently in interneurons. This impairs local autocrine signaling through the IGF-1 growth pathway, which impairs interneuron growth, leading to a reduction in the number of GABAergic synapses. We hypothesize that diminished synaptic inhibition contributes to the generation of epileptic spasms. In terms of spasms resolution, we propose that treatment with (1–3)IGF-1 restores IGF-1 levels in the cortex and re-establishes IGF-1R molecular signaling. This reinstates GABAergic synapse formation, restores synaptic inhibition, and abolishes spasms.

**Figure 12.**
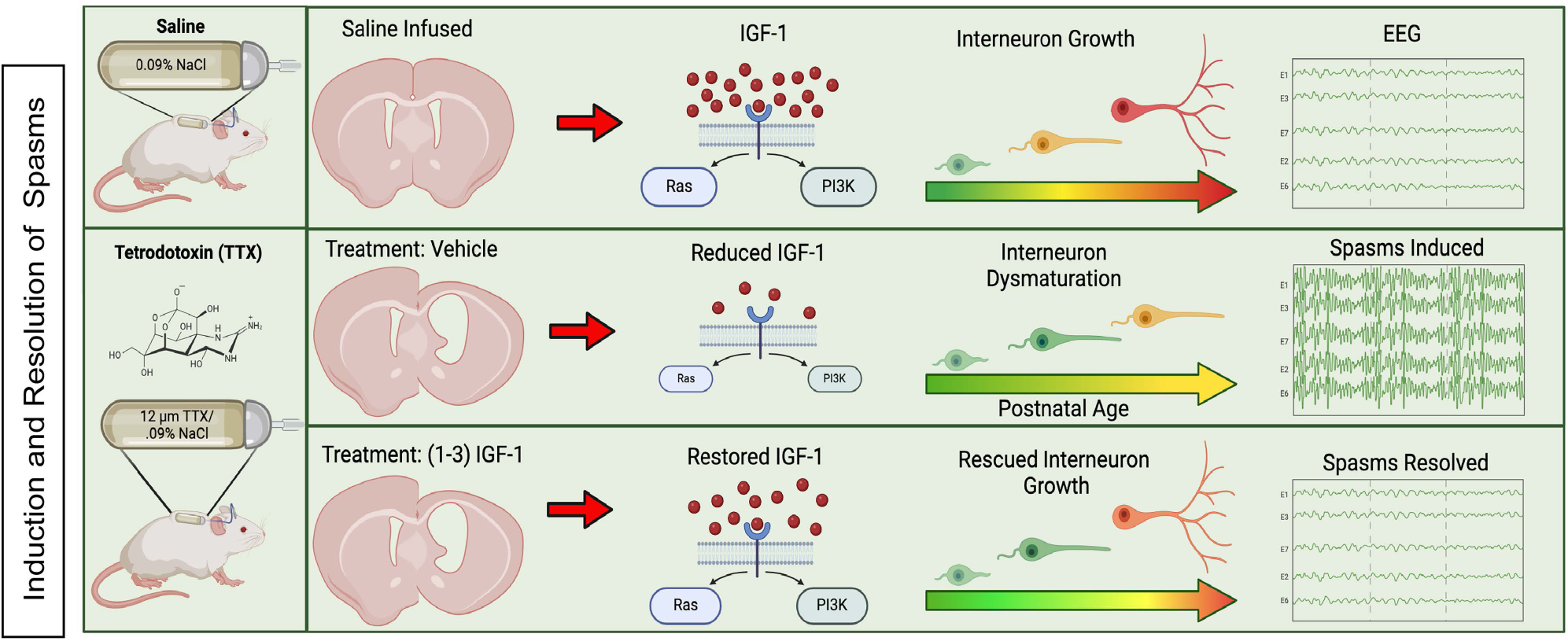
Hypothetical schemes for the role IGF-1 plays in the induction and resolution of epileptic spasms. In saline-infused control rats, IGF-1 signaling through the IGF-1R signaling pathways plays a role in interneuron growth. TTX-induced neocortical injury diminishes IGF-1 expression and reduces signaling through its growth pathway, resulting in interneuron dysmaturation (impaired maturation) that likely contributes to the induction of epileptic spasms. Treatment with (1–3)IGF-1 restores IGF-1 levels in the cortex, which activates its growth signaling pathway, rescues interneuron growth, and abolishes epileptic spasms. Created with BioRender.com.

We are not the first to implicate disruptions of inhibitory interneurons in the generation of epileptic spasms (see Introduction), but we are the first to suggest that interneuron dysmaturation can contribute to spasm generation. That said, dysmaturation may not arise from all etiologies of spasms and may be a unique consequence of early-life brain injury.

### Clinical Implications

Our examination of the neocortices of IESS patients following perinatal strokes indicates that the density of GABAergic nerve terminals is diminished (Figure 6) and IGF-1 levels are suppressed. These results are consistent with reports of reduced GABA and IGF-1 concentration in CSF taken from IESS symptomatic patients (23, 24). Taken together, these findings lead us to suspect that neocortical interneurons in patients with acquired structural brain abnormalities may be in a state of dysmaturation. If so, treatment with (1–3)IGF-1 or an analogue (67) may be one way to reinstate interneuron growth. If successful, it would be a novel disease-modifying therapy.

## Acknowledgements

This work was funded by CURE’s Infantile Spasms Initiative and NIH NINDS grants: RO1 NS105913 and R61/R33 NS112553 to J.W.S. It was also supported by IDDRC grant 1U54 HD083092 from the Eunice Kennedy Shriver National Institute of Child Health and Human Development. We thank the Neuropathology and Molecular Neuropathology Laboratories at Texas Children’s Hospital (TCH) for their assistance with human studies. The authors would like to acknowledge: Dr. Dinghui Yu and the IDDRC Microscopy Core at the NRI; the Small Animal Imaging Facility at TCH and its director’s assistant, Dr. Robia Paulter; and Dr. Mingshan Xue for his advice on AAV transfections.

## Author Contributions

C.J.B-R. and J.W.S. contributed to the conception and design of the study; C.J.B-R., J.T.L., T.L., A.E.A., J.D.F. Jr., and J.W.S. contributed to the acquisition and analysis of data; C.J.B-R., J.T.L., J.D.F. Jr., and J.W.S. contributed to drafting the text and preparing the figures.

## Declaration of Interests

J.W.S., J.T.L., and J.D.F. Jr. are inventors in a patent (US Patent 11,351,229) held by Baylor College of Medicine that covers therapies for treating infantile spasms and other treatment-resistant epilepsies.

---

**Supplemental Figure 1.**
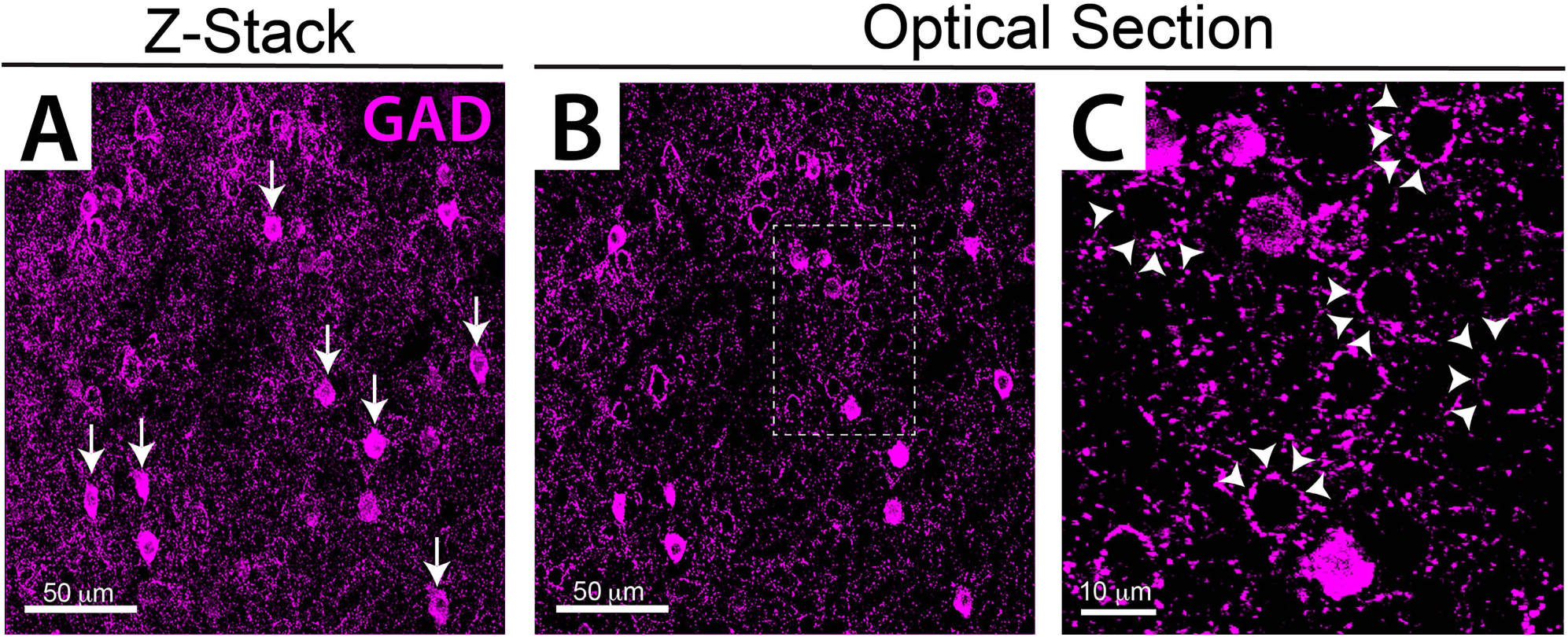
Dense networks of neocortical GABAergic nerve terminals revealed by GAD immunohistochemistry. (A) A stack of 14 confocal slices covering a depth of 10.5 µm that in addition to illustrating numerous GAD-positive cell bodies (arrows) displays a complex labyrinth of putative presynaptic GABAergic nerve terminals. (B) A single optical section taken from the images in A that further illustrates the complex microanatomy. Box outlined with dashed line is shown at higher magnification in C. (C) Innumerable GAD-positive presynaptic nerve terminals are shown as well as putative perisomatic basket cell like structures (arrow heads). Images were taken from neocortical layer V.

**Supplemental Figure 2.**
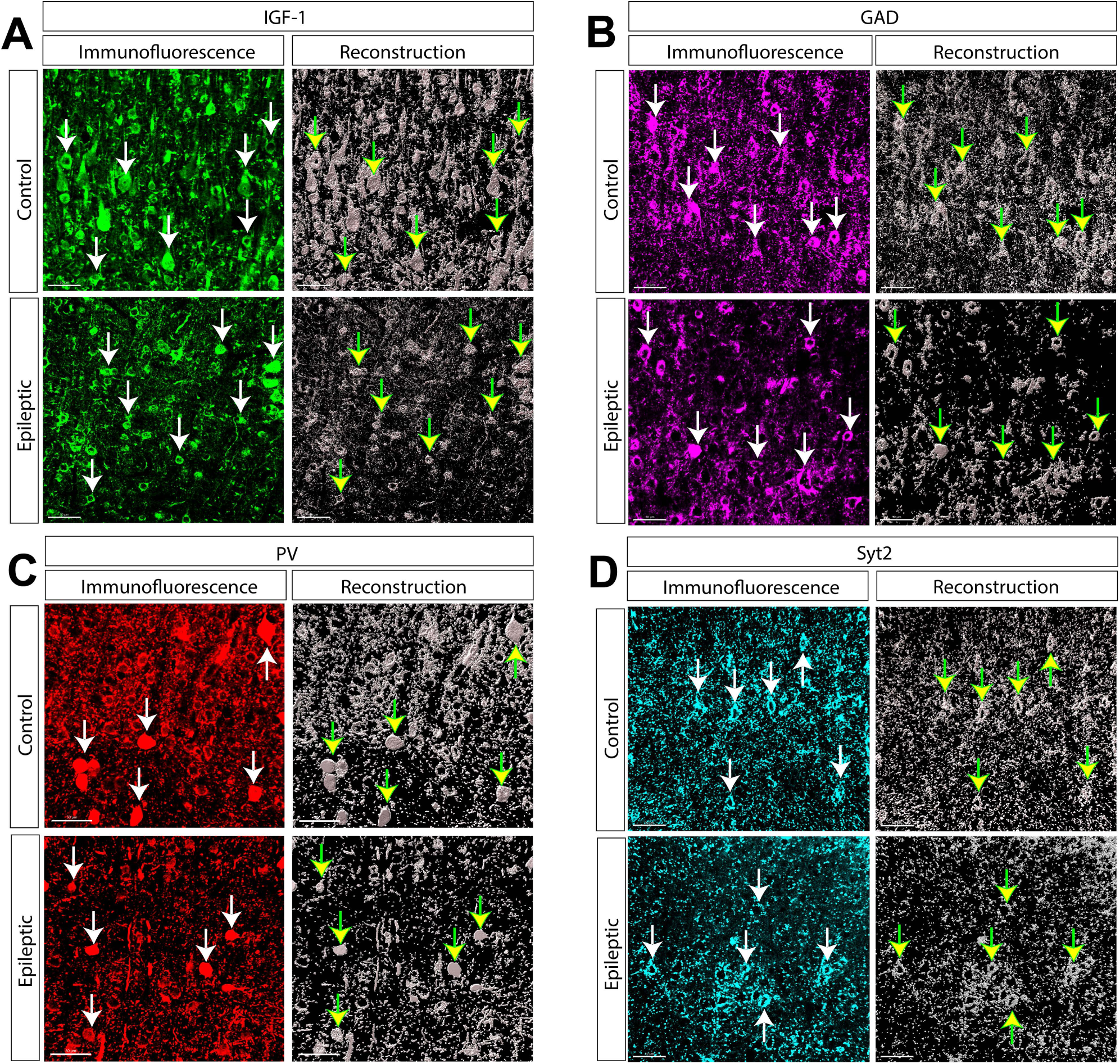
Representative examples of the differences in expression of IGF-1, GAD, PV and synaptotagmin 2 between control and experimental animals as delineated by 3D surface rendering and reconstruction of immunolabeling. Such reconstructions permitted the quantification of the neocortical volume occupied by immunoreactive cellular elements as reported in Figure 4C and D. (A) IGF-1, (B) GAD, (C) PV and (D) synaptotagmin 2. In each panel, confocal images on the left (in color) show immunohistochemistry for the molecule. On the right are the reconstructions shown as monochrome. Arrows denote the same cell bodies in immunohistochemical staining and reconstructions. In the case of synaptotagmin 2, the cell bodies denoted are enveloped by a dense plexus of synaptotagmin 2 immunoreactive nerve terminals. All images are from layer V of neocortical area S1 ipsilateral to the TTX-induced lesion.

**Supplemental Figure 3.**
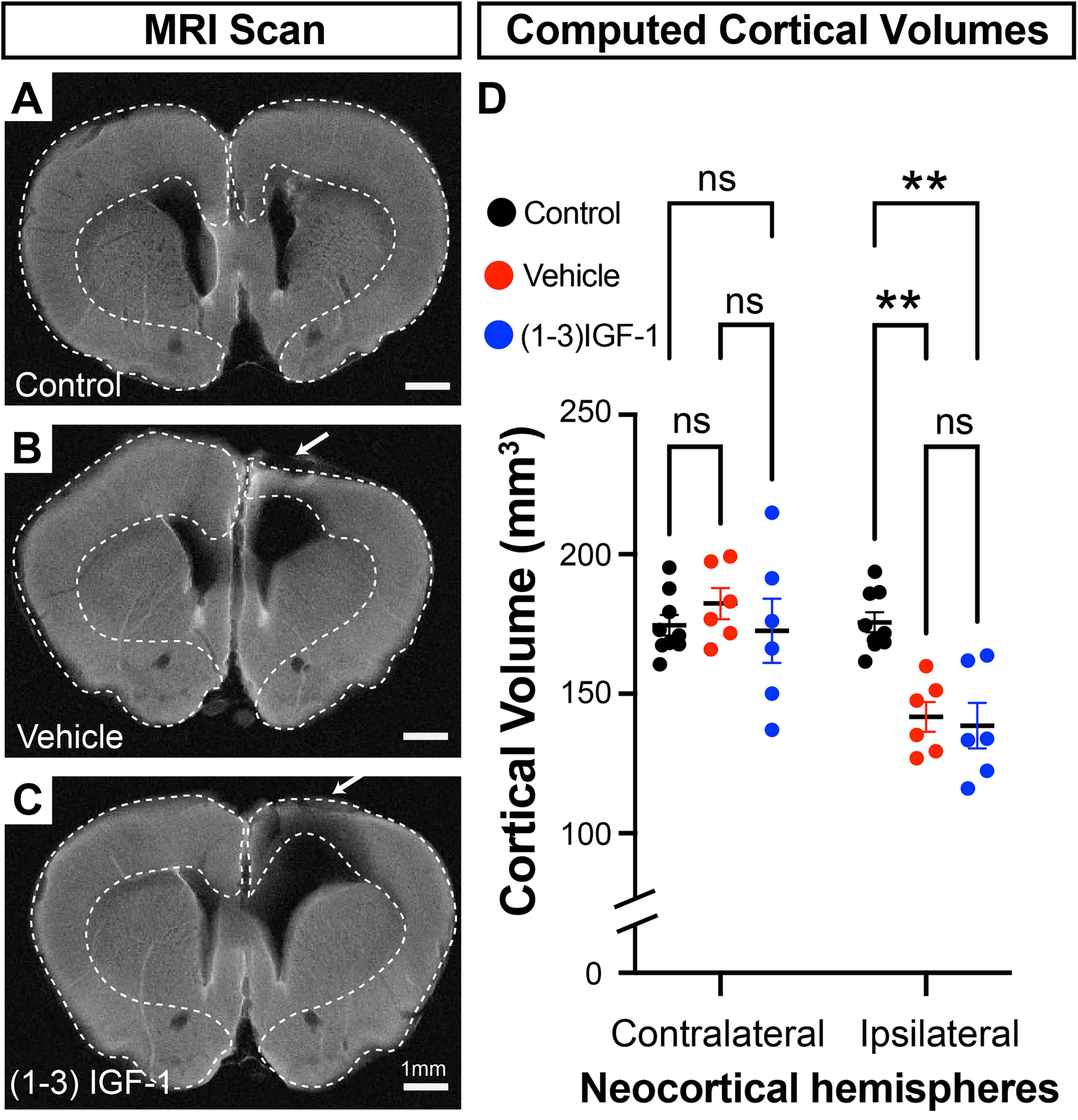
Treatment with (1–3)IGF-1 did not alter the volume of TTX-induced neocortical lesions. (A-C) MRI slices taken from a representative animal from the three treatment groups. Arrows denote the TTX-induced lesions. Dashed lines outline the perimeter of the neocortex that was used in all MRI slices from each animal to compute total neocortical volume. (D) The computed cortical volumes showed that infusion of TTX did not alter the volume of the cortex contralateral to TTX infusion. However, in both the vehicle and (1–3)IGF-1 treated rats, cortical volumes were reduced compared to control animals, However, no difference was detected in the neocortical volumes between the vehicle and (1–3)IGF-1 treated animals indicating that treatment with the tripeptide did not reduce or increase the size of TTX-induced lesion. Ipsilateral Vehicle: 141.8 ± 5.4 mm^3^ vs Ipsilateral (1–3)IGF-1: 138.6 ± 8.2 mm^3^ p = 0.999 - two-way ANOVA with correction for multiple com

**Supplementary Table 1.** Summary of Resected Surgery Patients’ Neuropathological Findings, Seizure Classification and Type of Surgery Undertaken. Patient ID: S-IESS patient number, C-Tumor control patient number.

| ID | SEX | PATHOLOGY | SEIZURE (SZ) TYPE | AGE AT SURGERY (YRS) | SURGERY TYPE |
| --- | --- | --- | --- | --- | --- |
| S1 | F | INTRAUTERINE STROKE, INTRAVENTRICULAR HEMORRHAGE, REFRACTORY EPILEPSY WITH CLUSTER SEIZURES | INFANTILE SPASMS AT 6 MONTHS OF AGE - LATER DEVELOPED FOCAL MOTOR SZ | 3 | RIGHT HEMISPHERECTOMY |
| S2 | F | DOWN SYNDROME, PERINATAL STROKE, REFRACTORY INFANTILE SPASMS | INFANTILE SPASMS AT 6 MONTHS SPASMS PERSISTED UNTIL SURGERY | 3 | LEFT HEMISPHERECTOMY |
| S3 | M | GRADE 4 LEFT INTRAVENTRICULAR HEMORRHAGE REFACTORY EPILEPSY | INFANTILE SPASMS AT 8 MONTHS - LATER DEVELOPED FOCAL MOTOR SZ | 1.75 | LEFT HEMISPHERECTOMY |
| S4 | M | VEIN OF GALEN MALFORMATION AND RIGHT-HEMISPHERIC ARTERIOVENOUS MALFORMATION, REFRACTORY SPASMS | INFANTILE SPASMS AT 6 MONTHS; SPASMS PERSISTED UNTIL SUGERY | 4 | RIGHT HEMISPHERECTOMY |
| C1 | M | GLIOMA WITH MIXED OLIGODENROGLIOMA, ANGIOCENTRIC GLIOMA AND DYSEMBRYOPLASTIC NEUROEPITHELIAL TUMOR | ALTERED AWARENESS, FOCAL MOTOR SZ PRIOR AND LEADING TO TUMOR DIAGNOSIS | 6 | LEFT ANTERIOR TEMPORAL GLIOMA RESECTION |
| C2 | F | ANGIOCENTRIC GLIOMA, | FOCAL MOTOR SZ,<br>PRIOR AND<br>LEADING TO<br>TUMOR<br>DIAGNOSIS | 8 | LEFT TEMPORAL<br>ANGIOCENTRIC<br>GLIOMA RESECTION |
| C3 | F | DYSEMBRYOPLASTIC<br>NEUROEPITHELIAL<br>TUMOR (DNET) | FOCAL MOTOR SZ,<br>PRIOR AND<br>LEADING TO<br>TUMOR<br>DIAGNOSIS | 5 | RIGHT ANTERIOR<br>MEDIAL TEMPORAL<br>DNET RESECTION |
| C4 | M | GANGLIOGLIOMA, | FOCAL MOTOR SZ,<br>PRIOR AND<br>LEADING TO<br>TUMOR<br>DIAGNOSIS | 12 | RIGHT ANTERIOR<br>TEMPORAL<br>GANGLIOGLIOMA<br>RESECTION |

## Notes

### Competing Interest Statement

The authors have declared no competing interest.

